# A genome-wide Approximate Bayesian Computation approach suggests only limited numbers of soft sweeps in humans over the last 100,000 years

**DOI:** 10.1101/2019.12.22.886234

**Authors:** Guillaume Laval, Etienne Patin, Pierre Boutillier, Lluis Quintana-Murci

## Abstract

Over the last 100,000 years, humans have spread across the globe and encountered a highly diverse set of environments to which they have had to adapt. Genome-wide scans of selection are powerful to detect selective sweeps. However, because of unknown fractions of undetected sweeps and false discoveries, the numbers of detected sweeps often poorly reflect actual numbers of selective sweeps in populations. The thousands of soft sweeps on standing variation recently evidenced in humans have also been interpreted as a majority of mis-classified neutral regions. In such a context, the extent of human adaptation remains little understood. We present a new rationale to estimate these actual numbers of sweeps expected over the last 100,000 years (denoted by *X*) from genome-wide population data, both considering hard sweeps and selective sweeps on standing variation. We implemented an approximate Bayesian computation framework and showed, based on computer simulations, that such a method can properly estimate *X*. We then jointly estimated the number of selective sweeps, their mean intensity and age in several 1000G African, European and Asian populations. Our estimations of *X*, found weakly sensitive to demographic misspecifications, revealed very limited numbers of sweeps regardless the frequency of the selected alleles at the onset of selection and the completion of sweeps. We estimated ∼80 sweeps in average across fifteen 1000G populations when assuming incomplete sweeps only and ∼140 selective sweeps in non-African populations when incorporating complete sweeps in our simulations. The method proposed may help to address controversies on the number of selective sweeps in populations, guiding further genome-wide investigations of recent positive selection.

## Introduction

Evaluating the legacy of positive, Darwinian selection in the human genome has proved crucial for identifying the genes underlying the broad morphological and physiological diversity observed across human populations, and for increasing our understanding of the genetic architecture of adaptive phenotypes. Genome-wide scans for positive selection have been largely guided by the selective sweep model (Hernandez, et al. 2011), a model in which advantageous mutations increase in frequency until fixation under the pressure of positive selection (Maynard Smith and Haigh 1974), see (Pritchard, et al. 2010) for a review. These studies have provided a flurry of candidate loci, e.g., (Voight, et al. 2006; Frazer, et al. 2007; Sabeti, et al. 2007; Tang, et al. 2007; Williamson, et al. 2007; Pickrell, et al. 2009b; Grossman, et al. 2010; Granka, et al. 2012; Grossman, et al. 2013; Fagny, et al. 2014; Pybus, et al. 2015; Schrider and Kern 2017), see.(Vitti, et al. 2013; Jeong and Di Rienzo 2014; Fan, et al. 2016) for reviews.

There is a little overlap across different genome-wide studies and false discovery rates remain high (Teshima, et al. 2006; Akey 2009; Hsieh, et al. 2016). Nevertheless, the lowest empirical *P*-values (*P*-values computed from genome-wide observed data) tend to remain significant after the computation of *P*-values from neutral computer simulations (Hsieh, et al. 2016), suggesting that top candidates usually put forward in genome-wide scans contain a fraction of true positives. The top candidates detected in the 1000 Genomes (1000G) CEU population (Auton, et al. 2015) using simulated *P*-values contain ∼5 genomic regions with selection signals found significant after correction for the high numbers of tests performed genome-wide (-log(*P*-value)>8) (Grossman, et al. 2013). Unfortunately, it is fairly well-known that such multiple testing corrections applied to discard false positives also discard true targets of selection, because increasing detection thresholds also reduce the statistical power to detect sweeps (Pavlidis and Alachiotis 2017).

Recently, simulation-based machine learning algorithms implemented to detect selective sweeps (Pybus, et al. 2015; Schrider and Kern 2016, 2017) and classify them into hard or soft sweeps, revealed up to ∼1,000 soft sweeps in non-African 1000G populations (Schrider and Kern 2017). The hard sweeps refer to sweeps on *de novo* advantageous mutations (frequency equal to 1/2*N* at the onset of selection) (Maynard Smith and Haigh 1974; Hermisson and Pennings 2005). In contrast, the soft sweep model as investigated by (Schrider and Kern 2017) refers to selective sweeps on standing variation as defined in (Hermisson and Pennings 2005), i.e. mutations at frequency higher than 1/2*N* at the onset of selection (Orr and Betancourt 2001; Innan and Kim 2004; Hermisson and Pennings 2005; Przeworski, et al. 2005; Pennings and Hermisson 2006a).

However, Harris and colleagues (2018) argued that these high numbers of soft sweeps can be predicted on the basis of the error rates observed when classifying neutral sites, highlighting a spurious inflation of the numbers of detected soft sweeps due to false positives (see (Harris, et al. 2018) for details). This recent debate illustrates the lack of *ad-hoc* methods that formally estimate actual numbers of selective sweeps in populations (e.g., numbers of sweeps occurred over a given time span). Genome-wide scans for selection are designed to detect sweeps but the numbers of detected/classified sites, e.g., (Li and Stephan 2006; Pybus, et al. 2015; Schrider and Kern 2017), poorly estimate the actual numbers of sweeps because of the burden of false positives. When using weakly or moderately stringent detection thresholds, the lists of detected sweeps are potentially enriched in false positives (the majority of detected sweeps can be neutral regions with spurious signal of selection). Therefore, as mentioned above, in genome-wide studies we are forced to apply stringent detection thresholds to discard these false positives. This may drastically reduce the number of detected sweeps (see the ∼862 and ∼18 sweeps on average retained with the default and a most stringent probability threshold of 0.9 used in the machine learning classifier (Schrider and Kern 2017; Harris, et al. 2018)). These reduced numbers are likely far from being representative of all true targets of selection. Indeed, many true targets of selection cannot be detected because of a diminished statistical power when applying stringent thresholds, resulting in an unknown fraction of false negatives, e.g., sweeps of weak intensity (weak selection coefficients or very recent ages of selection) already hardly detectable even at nominal thresholds of P<0.01. As a consequence, despite the high number of genome-wide selection scans performed and the development of sophisticated methods, the numbers of loci truly under positive selection in populations and, therefore, the contribution of positive selection to recent human adaptation remain poorly understood, from rare numbers of classic sweeps (Pritchard, et al. 2010; Hernandez, et al. 2011) to high numbers of soft sweeps (Schrider and Kern 2017).

Overall, current methods are therefore related to numbers of detected sweeps not to actual numbers of selective sweeps in populations, which can markedly differ because of reasons described above (statistical power and detection thresholds). In this manuscript, we present a simulation-based method that directly estimates these actual numbers of selective sweeps in populations, denoted by *X* for the rest of the study. We did not attempt to detect sweeps separately. We rather considered *X* as a parameter in a model of the genome-wide effects of positive selection. To estimate *X*, we implemented an approximate Bayesian computation (ABC) method (Beaumont, et al. 2002) based on summary statistics that summarize the genome-wide signals of selection and correlate with *X*, as shown below by computer simulations (see the results section). The ABC method presented here does not output a list of detected sweeps but can help to design and/or interpret normal genome-wide scans by providing general indications about the numbers of sweeps really expected in the analyzed populations (scripts and software are available on demand).

We applied this method to several African, European and Asian 1000G populations (Online Methods, Supplementary Table S1). In particular, we aimed to show the suitability of our approach in the context of the high numbers of sweeps previously found, by considering hard and soft sweep models as investigated by Schrider and Kern (Schrider and Kern 2017). We thus considered selection models whereby advantageous mutations exhibit a non-zero selection coefficient *s* since a specific time *t*. We considered *t* ranged from present to 3,500 generations ago (the last 100,000 years assuming a generation time of 28 years (Fenner 2005; Moorjani, et al. 2016)}) and frequency of the selected allele when the sweep begins, denoted by *p_start_*, ranged from 1/2*N* (hard sweeps) to 0.2. This corresponds to the same soft sweep scenario as investigated by Schrider and Kern (Schrider and Kern 2017), who ignored the soft sweep scenario that considers multiple independent advantageous mutations at a single locus (Garud, et al. 2015). Note that our simulation-based method can easily be updated in order to also incorporate soft sweeps on multiple advantageous mutations. Finally, we considered lower selection coefficient values, i.e., *s* ranged from 0.001 to 0.05, and avoided using flat distributions as previously done (Schrider and Kern 2017). We rather randomly drawn *s* from distributions enriched in low values as expected (e.g., ∼60% of mutations were predicted nearly neutral or with a moderate effect on fitness, i.e., with *s*≤1% (Boyko, et al. 2008)).

We jointly estimated *X* and the numbers of sweeps with very low (1/2*N*≤*p_start_*<0.01), low (0.01≤*p_start_*<0.1) and intermediate (0.1≤*p_start_*<0.2) initial frequencies of the selected alleles, denoted by *X*_1_*, X*_2_ and *X*_3_ respectively (in our model X = X_1_ + X_2_ + X_3_). We did not estimate the number of hard sweeps *stricto senso* (*p_start_*=1//2*N*) as classically performed (Schrider and Kern 2017). Instead, we merged hard and soft sweeps with very low *p_start_* into the same category *X*_1_ because such sweeps left genomic signatures virtually indistinguishable (see (Jensen 2014) for a review). For example, the probability to misclassify soft sweeps as hard sweeps increases when the initial frequency of the selected allele approaches 1/2*N* (Peter, et al. 2012) since at such low frequency these two scenario generate similar signals of selection (Przeworski, et al. 2005; Ferrer-Admetlla, et al. 2014; Jensen 2014). Finally, we jointly estimated the *X*s (*X*, *X*_1_*, X*_2_ and *X*_3_) and the average intensity and age of selection, 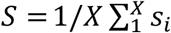 and 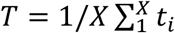, with *s*_i_ and *t*_i_ being the intensity and age of the *i*^th^ selective sweep.

## Results

### Overview of the ABC Approach

For the clarity of the manuscript, we present in this section an overview of the rational and methods implemented in our ABC approach.

ABC approaches are based on simulated data that mimic the empirical data to analyze and on statistics that both summarize each dataset and exhibit a monotonic relationship with the parameter to estimate. Here we simulated whole-genome sequence (WGS) data that mimic the 1000G data, i.e., ∼3Gb of DNA sequences for about 100 individuals sampled per population. We used summary statistics based on the widely used comparison between neutral mutations, labelled here ENVs (Evolutionary Neutral Variants), and mutations potentially targeted by selection, labelled PSV (Possibly Selected Variants), e.g., synonymous and non-synonymous comparison (Kimura 1977; Mcdonald and Kreitman 1991). Our rational was motivated by previous studies showing that positive selection causes genome-wide excesses of candidate SNPs of selection (i.e., those with extreme values for a given neutrality statistic) within or nearby genes relative to intergenic regions (e.g., coding SNPs or *cis*-acting eQTLs *vs* intergenic SNPs (Voight, et al. 2006; Frazer, et al. 2007; Barreiro, et al. 2008; Kudaravalli, et al. 2009; Jin, et al. 2012; Fagny, et al. 2014; Schmidt, et al. 2019)). In such studies, the ENV SNP class is the neutral baseline used to control for false discoveries (similar rates of false positive are expected across SNP classes (Barreiro, et al. 2008)). As summary statistic, we used the odds ratio for selection (OR) (Kudaravalli, et al. 2009), which assesses this genome-wide excess of candidate SNPs in PSVs relative to ENVs using all candidate SNPs treated equally regardless of their individual *P*-values. The OR being approximately a ratio between the percentages of candidate SNPs in PSVs and ENV, this summary statistic is thus expected close to one under neutrality and above one under selection (Kudaravalli, et al. 2009; Fagny, et al. 2014). Based on computer simulations, we show in the next section that the OR exhibits a monotonic relationship with *X*.

The ENVs are easy to conceptualize. In real data, they are intergenic SNPs at some distance from the nearest gene and purged from any functional sites to be unaffected by selection. In contrast, the PSVs contains all potential targets of selection and neutral nearby SNPs (Supplementary Fig. S1). For example, in real data the PSVs correspond to nonsynonymous mutations and their neighbors (e.g., synonymous and intronic mutations) or regulatory mutations and their neighbors (e.g., variants located upstream/downstream of genes or in remote regulatory regions). We need to incorporate in PSVs these nearby SNPs in order to take into account the hitchhiking effects of positive selection. Indeed, selective sweeps produce clusters of candidate SNPs in the vicinity of the targets of selection whereas under neutrality candidate SNPs are more uniformly scattered (Voight, et al. 2006). The magnitude of such clustering depends on the intensity and age of selection and thus provides information for our estimations of *X*, *S* and *T*. In simulations, the PSVs are randomly defined (the remaining mutations are ENVs) and contain *X* selective sweeps of various intensity, age and frequency of the selected alleles at the onset of selection (each PSV region does not contained a target of selection but all simulated targets of selection are defined as PSVs).

The desired ABC posterior distributions are computed in each 1000G population by comparing simulated and empirical WGS datasets each summarized by a vector of *K* ORs, *K* being the number of neutrality statistic used. Because background selection (BGS) may generate spurious positive selection signatures (Coop, Pickrell, Novembre, et al. 2009; Pritchard, et al. 2010; Hernandez, et al. 2011), we decided to focus on neutrality statistics previously found insensitive to BGS (Zeng, et al. 2006; Fagny, et al. 2014), i.e., Fay and Wu’s *H* (F&W-*H*) (Fay and Wu 2000), iHS (Voight, et al. 2006), DIND (Barreiro, et al. 2009), ΔiHH (Grossman, et al. 2010), and XP-EHH (Sabeti, et al. 2007). Overall, the posterior distributions obtained are the parameter values underlying the simulated ORs that best fit with the empirical 1000G ORs. We use posterior mean as point estimate and the 95% credible intervals (CI) boundaries to assess the uncertainty of the estimations.

### Odds Ratio for selection reflects the number of selective sweeps

To test the unknown relationships between *X* and the OR for selection, we simulated WGS data with different fixed numbers of selective sweeps, X = 0, 50, 100,150] (Online Methods). The simulated WGS were obtained using recombination profiles randomly drawn from human recombination maps (Frazer, et al. 2007) and a demographic model previously inferred, as classically done (Peter, et al. 2012; Grossman, et al. 2013; Nakagome, et al. 2016; Schrider and Kern 2017; Smith, et al. 2018; Uricchio, et al. 2019) (Online Methods). Because we aimed to gain information by incorporating inter-population statistics such as the XP-EHH, we did not use one-population models, as used by Schrider and Kern (Schrider and Kern 2017). Instead, we used a three-populations demographic model calibrated to replicate the allele frequency spectrum, population subdivision and linkage disequilibrium in the YRI (Africa), CEU (Europe) and CHB (Asia) populations (Schaffner, et al. 2005; Grossman, et al. 2010; Grossman, et al. 2013) and used by (Grossman, et al. 2013; Pybus, et al. 2015) to detect selective sweeps in the 1000G populations. Specifically, to simulate the *X* selective sweeps in one population we assumed neutrality (*X*=0) in the two other populations, e.g., to simulate *X* sweeps in Africa (YRI) we jointly simulated the European (CEU) and Asian (CHB) populations as two neutral references populations. We incorporated the *X* selected regions within simulated neutral DNA sequences to form ∼3Gb of WGS in which the randomly defined ENVs and PSVs respectively cover 30% and 70% of the genome (these proportions corresponds to the proportions used to analyze the 1000G populations, see below). Finally, for each neutrality statistic used we defined candidate SNPs (top 1% of SNPs) (Online Methods) and we computed the ORs (Online Methods) separately for the F&W-H, iHS, DIND, ΔiHH, and for two XP-EHHs (we obtained a total of 6 ORs since in a three-populations branching model, as the one used in this study, there are two XP-EHH per population).

To simulate *X* sweeps per WGS data (Online Methods), *p_start_* was randomly drawn from the allele frequency spectrum at the generation *t* excluding values higher than 0.2 in agreement with (Schrider and Kern 2017). The *s* and *t* parameters were randomly drawn from flat distributions, s ∼ U(0.001, 0.05) and t ∼ U(0, 100) *kya*, and we excluded complete sweeps (sweeps at fixation) using a rejection algorithm. By excluding complete sweeps, we reproduced in our simulations the excess of mutations with low or moderate effect on fitness previously evidenced (Boyko, et al. 2008). Indeed, the resulting distributions of *s* (and also *t*) really used for our simulations are enriched in low values (Fig. 1A), since (all other things being equal) complete sweeps tend to be stronger (or older) than incomplete sweeps. However, we started our analysis by excluding complete sweeps because previous results support limited numbers of complete sweeps in humans, e.g., lack of extreme differences in allele frequency between populations due to fixation events (Coop, Pickrell, Kudaravalli, et al. 2009; Pritchard, et al. 2010; Hernandez, et al. 2011). This data-driven assumption was further relaxed by considering both complete and incomplete sweeps in our simulations (see the sections below).

**Fig. 1.**
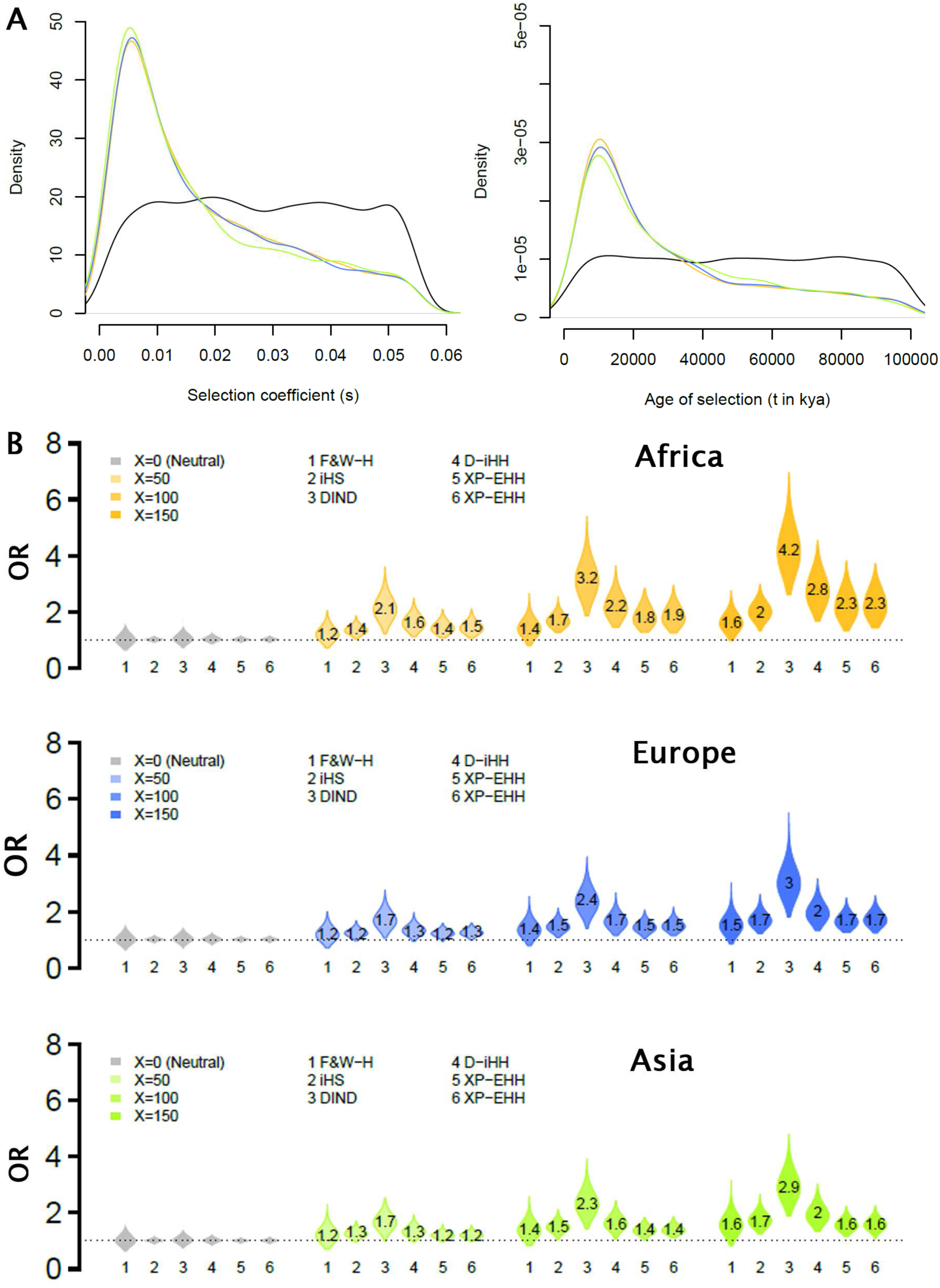
Odds ratio for selection mirror the number of selective sweeps in simulated populations. Relationship between the odds ratio for selection (OR) and the number of selective sweeps, in computer simulations of African, European and Asian populations, using various numbers of sweeps, X = 0, 50, 100, 150] (1,000 simulated WGSs for each). Simulated PSVs are randomly defined and cover 70% of the genome, a value corresponding to the percent of PSVs defined in the 1000G population. The *X* selective sweeps were simulated in the African, European or an Asian population, using the two other populations as neutral reference for interpopulation statistics. Each selective sweep has been simulated by considering frequency at the onset of selection varying from 1⁄2N to 0.2 and randomly drawn from the allele frequency spectrum at the onset of selection. For each selective sweep the intensity and the age of selection were randomly drawn from uniform distributions, s ∼ U(0.001, 0.05) and t ∼ U(0, 100) kya. We excluded complete sweeps by only keeping simulated sweeps with current selected allele frequencies ranged from 0.2 to 0.95. (**A**) Distributions of the *s* (left hand side) and *t* (right hand side) used to simulate the *X* incomplete sweeps in the African (yellow), European (blue) and Asian (green) population. The initial uniform distributions are indicated in black. (**B**) Simulated ORs for Fay & Wu’s *H* (F&W-*H*), iHS, DIND, ΔiHH (D-iHH) and two pairwise XP-EHH. The average of the simulated OR are reported within each violin plot.

We observed a monotonic relationship between *X* and the ORs, i.e., the six ORs increase with *X* in every simulated scenario (Fig. 1B, supplementary Fig. S2). As expected, the values of ORs depend on the frequencies of advantageous alleles at the onset of selection (supplementary Fig. S2). The lowest ORs were obtained when simulating sweeps with initial frequencies of the advantageous alleles, *p_start_*, ranged from 0.01 to 0.2. Indeed, soft sweeps typically have weaker effects on linked sites (Przeworski, et al. 2005; Pennings and Hermisson 2006b; Pritchard, et al. 2010), resulting in a low enrichment of candidate SNPs in the vicinity of the targets of selection. The highest simulated ORs were obtained for hard sweeps *stricto senso* (*p_start_*=1/2*N*) and soft sweeps with very low initial frequencies (1/*N≤p_start_*<0.01). In addition, simulated ORs obtained for these two scenario were found to be very similar, as expected (Przeworski, et al. 2005; Peter, et al. 2012; Ferrer-Admetlla, et al. 2014; Jensen 2014). This observation confirms that hard sweeps and soft sweeps on very rare standing variation are almost indistinguishable on the basis of their outcomes (supplementary Fig. S2), which justifies the merging of these two scenarios. In the following, we do not provide separate estimates of the numbers of hard sweeps and only provide estimations of numbers of sweeps on very rare standing variants as a whole (*p_start_*<1%, *X*_1_).

### Odds Ratio for selection are poorly sensitive to demographic assumptions

As expected, the ORs values follow the power to detect selection as previously assessed for iHS and XP-EHH assuming various demographic histories (highest, intermediate and lowest power in the African, European and Asian demography respectively (Pickrell, et al. 2009b)). For example, the European and Asians ORs found lower than in Africa (Fig. 1B) reflect a lower enrichment of candidate SNPs, in agreement with the reduced power to detect selection in bottlenecked populations relative to expanding populations (Pickrell, et al. 2009a; Huff, et al. 2010; Gunther and Schmid 2011; Grossman, et al. 2013; Fagny, et al. 2014). Interestingly, only little difference in the behavior of ORs was observed across populations simulated with contrasting demographic histories (i.e., African expansions *vs* Eurasian bottlenecks, Fig. 1B).

Moreover, the Asian and European simulated ORs are virtual identical albeit slightly lower in Asia (e.g., 2.3 *vs* 2.4 in average for iHS with *X*=100, Fig. 1B) whereas the Asian bottleneck is four times stronger than in Europe. These observations suggest that the subsequent ABC estimations of *X* will be weakly affected by potential demographic misspecifications (this specific point is tested in the next sections below). Taken together, our results show that the OR for selection contain information on *X* (monotonic relationship, see above) without being largely affected by demography, which supports the use of such a summary statistic in an ABC framework.

### Accuracy of the ABC estimations

The ABC method was implemented using simulations identical to those presented in Fig. 1, except that *X* was randomly drawn in flat prior distributions, X ∼ U(0, 200) (Online Methods), opposed to fixed values (priors used for *s* and *t* are enriched in low values because we excluded complete sweeps, see Fig. 1A). To assess the accuracy of the estimations (Fig. 2) we simulated new WGSs following the same recombination and demographic model and treated them as empirical data for which the parameter values are known (Online Methods).

**Fig. 2.**
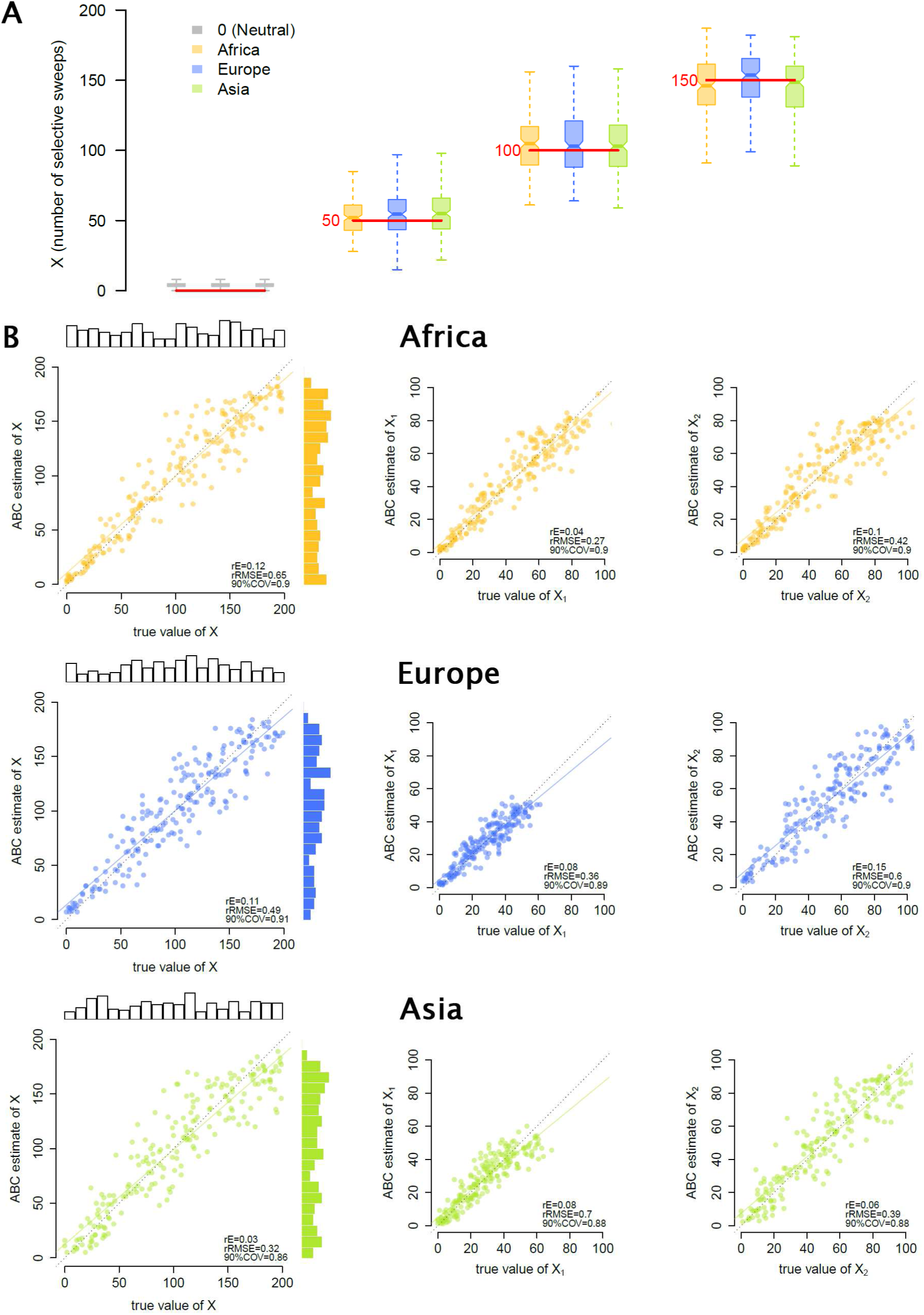
Accuracy of the ABC estimation of the numbers of selective sweeps. The estimations were performed using the six ORs shown in Fig. 1 and 105 ABC simulations obtained similarly as in Fig. 1 but using *X* randomly drawn in flat prior distributions, X ∼ U(0, 200). The priors used for *s* and *t* are shown in Fig. 1A. (**A**) ABC estimations of *X* performed in simulated WGSs used as empirical data containing 0, 50, 100 or 150 selective sweeps (200 simulated WGSs in each case). Horizontal red lines indicate the true simulated values of *X*. (**B**) Relationships between the ABC estimates (*y*-axis) and the corresponding parameter values (*x*-axis) obtained for the number of selective sweeps *X*, as well as for *X*<u>_1_</u> and *X*_2_, the numbers of sweeps with very low (1/2*N*≤*p_start_*<0.01) and low (0.01≤*p_start_*<0.1) frequencies of the selected allele at the onset of selection. The estimations of *X*_3_ (number of sweeps with intermediate initial frequencies, 0.1≤*p_start_*<0.2) are not shown because simply deduced from the relation, X = X_1_ + X_2_ + X_3_. In each case, the 200 simulated WGSs used as empirical data were generated following the same recombination and demographic model and using parameters drawn from the priors used for the estimations (top bar plots). Regression lines (colored lines) between true and estimated values and distributions of estimated parameters (right bar plots) are also indicated in each panel. Some classical accuracy indices are also reported, including the 90%*COV* (Online Methods). Additional information about the accuracy of the ABC estimations of *X*, *S* and *T* are shown in Supplementary Figs. S3-7 (the estimations of *S* and *T* corresponding to the panel **A** can be found in Supplementary Fig. S4).

We found unbiased estimations of *X* when simulating African, European or Asian demography (Fig. 2A), as shown by the linear correlations observed between ABC estimates and their corresponding true values (Fig. 2B). Such ABC estimates can thus be compared between populations since they are not biased by specific demographic events, such as the strong bottlenecks experienced by the non-African populations. We found a similar accuracy for the estimations of *X*_1_ (1/2*N*≤*p_start_*<0.01), *X*_2_ (0.01≤*p_start_*<0.1) and *X*_3_ (0.1≤*p_start_*<0.2) (Fig. 2B). We also estimated *S* and *T* in all the demographic scenario simulated (Supplementary Figs. S3-5) but found a diminished accuracy with respect to the estimations of *X* (e.g., underestimations and overestimations of *S* and *T* respectively when the selection is strong and recent in average, Supplementary Figs. S3-4). (As an internal control of the method, the CIs consistently predicted the range of values within which the true parameters were found (Supplementary Figs. S6-7, see also the 90%*COV* in Fig. 2B and Supplementary Figs. S3-4).

These results show that six ORs used in combination may successfully estimate *X*, with low or moderate credible intervals relative to the simulated priors (Supplementary Fig. S6). By analogy with normal genome-wide scans, our estimations of *X* seem to be weakly sensitive to false negative and positive signals. False positives do not contribute to ORs thanks to the use of ENVs as an internal neutral control, as illustrated by the estimations of *X* close to 0 under neutrality (Fig 2A) due to ORs fluctuating around one (i.e., similar rates of candidate SNPs across ENV and PSV classes, Fig 1B). In contrast, every selected region with more than 1% of candidate SNPs positively contributes to the OR proportionally to its intrinsic rate of candidate SNPs: selected regions with much more than 1% of candidate SNPs (e.g., true positives after multiple testing correction) contribute a lot and those with rates moderately higher than 1% (e.g., false negatives undetected after Bonferroni correction) also contribute but to a lesser extent. Moreover, thanks to the simulation of parameters that drive the statistical power to detect sweeps in normal selection scans (e.g., intensity of selection, demography), the model also predicts the expected fraction of sweeps with rates of candidate SNPs slightly below neutral thresholds (e.g., false negatives that cannot even be detected at unadjusted thresholds). Altogether, our estimations are not upward biased by neutral regions exhibiting spurious signatures of selection and also include a large fraction of sweeps that would not be detected by normal genome-wide scans (no systematic underestimation of *X*, Fig. 2).

However, these results have been obtained under a perfectly known demography while in real life, the demography is only partially known. Because our method, like every model-based method, should be affected by incorrect assumptions which may bias the estimations, we carefully tested the effect of demographic misspecifications on the estimations of *X* obtained in each 1000G population (see the next sections below).

### Limited numbers of incomplete selective sweeps in the 1000G populations

In this first round of estimations we assumed complete sweeps only. We thus analyzed 15 African, European and Asian 1000G populations separately (five populations per continent, Supplementary Table S1) using priors and simulations shown in Figs. 1A and 2B (the prior distribution used for *s* reproduces the expected enrichment of mutations with low or moderate effect on fitness (Boyko, et al. 2008)). As in simulations and for each population in which *X* was estimated, the two reference populations used to compute the interpopulation statistics were of differing continental origin, and were chosen from the populations previously used to calibrate the demographic model used (YRI, CEU or CHB) (Grossman, et al. 2010; Grossman, et al. 2013). Based on the VEP annotation of the 1000G data, we defined PSVs as all SNPs located within genic regions, including regions annotated as upstream and downstream which often contain regulatory elements, as well as any presumed regulatory sites located in intergenic regions (Supplementary Fig. S8, Online Methods). As already mentioned, variants neighboring functional mutations need to be included in PSVs to account for the clustering of candidate SNPs around putative targets of selection (Supplementary Fig. S1). Under this definition, PSVs account for ∼70% of the genome-wide variation (∼30% for ENVs).

We jointly estimated the *X*s (*X*, *X*_1_*, X*_2_ and *X*_3_), *S* and *T* in each population separately, using candidate SNPs defined genome-wide (top 1% of SNPs) and empirical ORs corrected for genomic variation in coverage, mutation and recombination rates, as recommended (Kudaravalli, et al. 2009) (Supplementary Fig. S9, Online Methods). In every population analyzed, the combination of these six empirical ORs used were found compatible with the simulated ORs outputted by the ABC model (Supplementary Fig. S10). We found a very limited numbers of selective sweeps (Fig. 3, Supplementary Fig. S11, see also the posterior predictive checks shown in Supplementary Fig. S12). For example, we estimated 65 [CI:37-96], 82 [CI:44-123] and 78 [CI:42-115] sweeps in the YRI, CEU and JPT (Japan) populations (Fig. 3A-C). For comparison, 810, 1,013 and 1,059 selective sweeps were previously found in these populations respectively (Schrider and Kern 2017). Overall, we estimated 62 [CI:36-91], 71 [CI:35-111] and 88 [CI:50-127] sweeps on average for the African, European and Asian continents (the estimates obtained in each population can be found in the Supplementary Table S2).

**Fig. 3.**
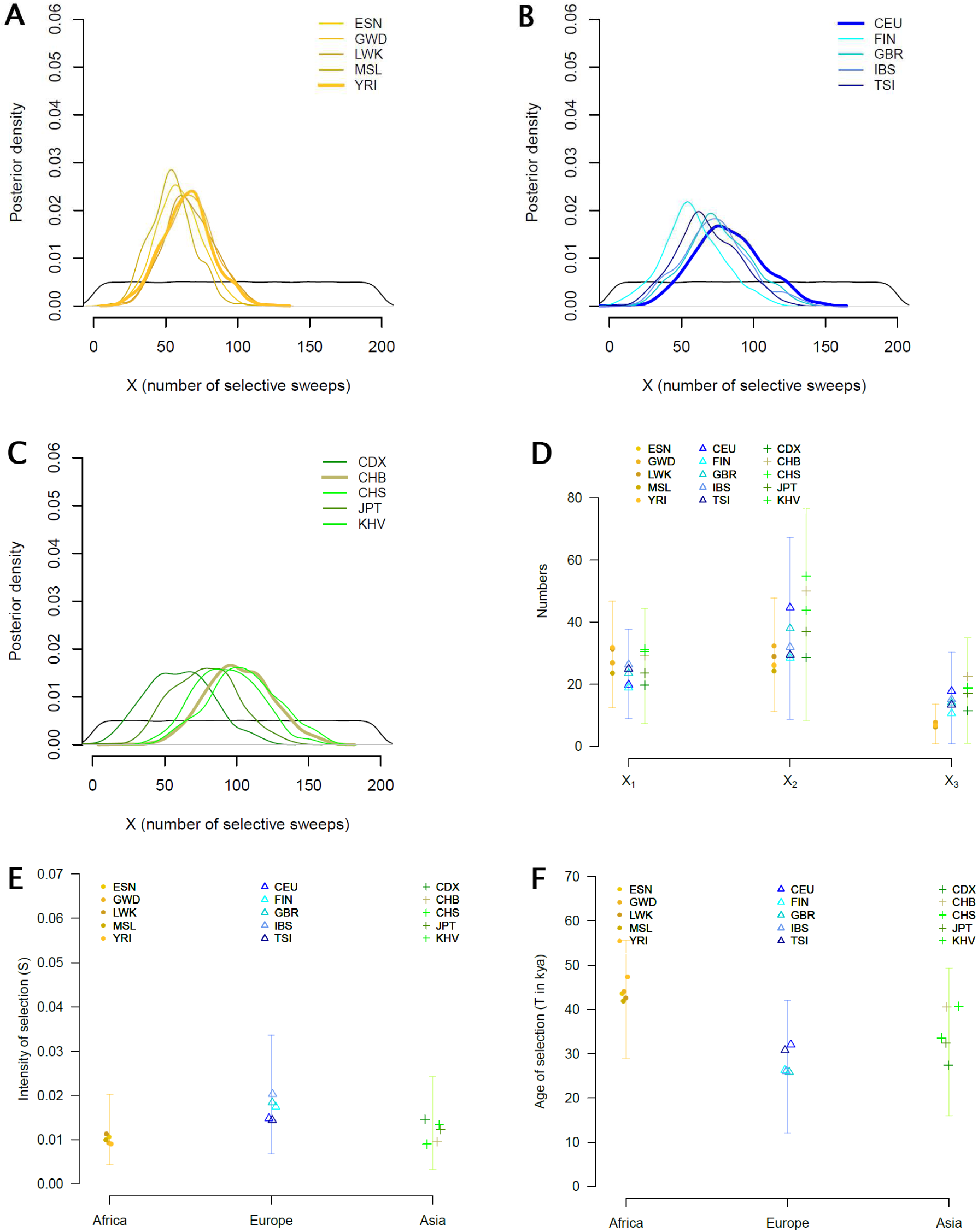
Estimations of the numbers of incomplete selective sweeps. The ABC estimations were performed in each 1000G population separately, using the same summary statistics and ABC simulations as used in Fig. 2B, i.e. six ORs, 10^5^ ABC simulations and *X* randomly drawn in flat prior distributions, X ∼ U(0, 200). The priors used for *s* and *t* are shown in Fig. 1A. The point estimates and CIs obtained in each population can be found in Supplementary Table S2. Posterior distributions of *X* in each (**B**) African, (**C**) European and (**D**) Asian population (priors are given in black). Populations used to calibrate the demographic model used (YRI, CEU and CHB) are indicated in bold. (**D**) ABC estimations of *X*_1_*, X*_2_ and *X*_3_, the numbers of selective sweeps with very low (1/2*N*≤*p_start_*<0.01), low (0.01≤*p_start_*<0.1) and intermediate (0.1≤*p_start_*<0.2) initial frequencies of the selected alleles. (**E,F**) ABC point estimates of the average of the intensity, *S*, and age of selection, *T*. The vertical bars shows the minimum and maximum estimates found across all populations considered.

### Robustness of the estimated numbers of selective sweeps

We checked the robustness of the estimations of *X* obtained in our first round of estimations shown in Fig. 3. First, we found consistent results using different combinations of summary statistics (Supplementary Fig. S13). We next checked the effects of potential demographic misspecifications (Supplementary Figs. S14-17). We adopted a strategy based on stress tests to evaluate the differences in numbers of estimated sweeps obtained when drastically modifying the demographic assumptions, e.g., replacing an expansion with a bottleneck or largely increasing bottleneck intensities (we used models in the range of demography previously inferred in human). We assessed this by swapping empirical and simulated ORs from differing continental regions, e.g., analyzing the African 1000G populations using ORs simulated under an Asian demographic model and inversely. Above all, we found very similar numbers of sweeps when setting such incorrect demographic scenarios in the model (Supplementary Figs. S15-16, details can be found in the Supplementary Table S2). The higher values of *X* estimated in Africa under an Asian demographic model, i.e., 85 [CI:51-121] (Supplementary Fig. S15) *vs* 62 [CI:36-91] sweeps on average, are simply due to lower simulated ORs (see the simulated ORs under African and Asian demography in Fig. 1B). The new estimations of *X* are increased because simulated ORs generated by larger *X* provide now a better fit with empirical ORs (see the graphical explanations given in Supplementary Fig. S14). We also re-analyzed the 1000G European populations under an Asian demographic model (Supplementary Fig. S16), i.e., now setting a bottleneck four times stronger than in our first round of estimations. The new estimations of *X* are only slightly increased, i.e., 83 [CI:48-120] (Supplementary Fig. S16) *vs* 71 [CI:35-111] sweeps on average, because ORs simulated under an Asian demography are only slightly lower than under an European demography (see also Fig. 1B and Supplementary Fig. S14). By symmetricity, we obtained smaller estimated *X* in Asia when assuming a European demography (Supplementary Figs. S14 and S16). Finally, we also modified several technical steps all at once, i.e., we set a 10% higher recombination rate in ENVs than in PSVs (Kong, et al. 2010), excluded selection signal found in ENVs (excluding ENVs in regions significantly enriched in candidate SNPs) and used empirical 1000G ORs computed merging all chromosomes together. We also found very similar, albeit slightly decreased, estimations of *X* (highly overlapping CIs when comparing with the first round of estimations, Supplementary Fig. S18).

Our stress tests confirmed the very limited numbers of sweeps found in this study. Notably, almost all re-estimated *X* fall within the 95% CIs boundaries of the initial estimations. Because a true demographic model does not exist, we did not used other models previously inferred in humans to re-estimate *X* (every inferred model poorly reproduces or totally ignores a given component of the human demography). We acknowledge that new estimations of *X* performed under other demographic models will differ but, given the low differences in numbers of sweeps found in our stress tests, they should not change in large proportions. What is of importance in the context of this study is that the very limited numbers of sweeps found cannot be explained by poorly inferred demographic parameters (see the little biases expected when assuming four time stronger/weaker bottlenecks than in reality, Supplementary Fig. S17).

### Limited numbers of sweeps when also considering complete sweeps

For reasons mentioned above, our first round of estimations was performed considering incomplete sweeps only whereas the high numbers of soft sweeps previously detected mainly refer to complete sweeps (Schrider and Kern 2017). We thus relaxed our assumption by simulating both complete and incomplete selective sweeps using flat prior distributions for *t*, as assumed previously (Schrider and Kern 2017), and several priors for *s*. The estimates obtained under these priors, including the flat prior for *s* previously used (Schrider and Kern 2017), are given in the Supplementary Fig. 19 and Supplementary Table S2. When considering priors enriched in small *s* values by using an equal mix between a Gamma distribution (60% of *s*≤0.01) (Boyko, et al. 2008) and a L-shape distribution (90% of *s*≤0.01), our method also provides unbiased estimations of *X* (Fig. 4A), since the ORs also reflect the *X* (Fig. 4B). As expected, the new estimated values of *X* are increased with respect to our first round of estimations, i.e., 115 [CI:68-160] and 165 [CI:119-211] sweeps on average in Europe and Asia (Fig. 4C, Supplementary Table S2), because of lower simulated ORs (Fig. 4B), in agreement with the reduced power to detect sweeps at fixation using neutrality statistics such as iHS (Voight, et al. 2006). However, our estimations are still far from the thousand of sweeps previously estimated per non-African population (Schrider and Kern 2017).

**Fig. 4.**
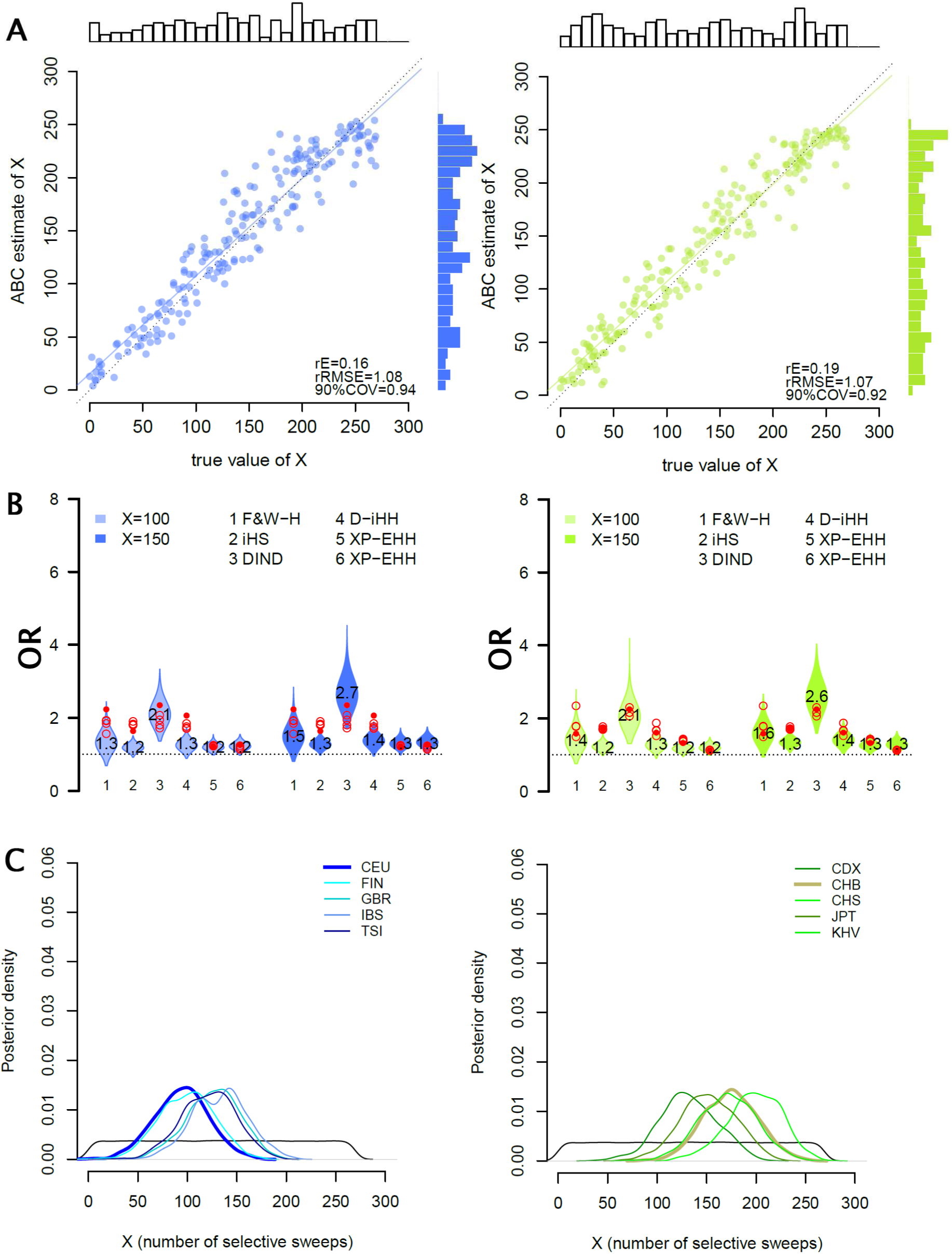
Estimations of the numbers of sweeps including complete selective sweeps. Simulations performed using the same recombination and demographic models used in Figs. 1-3 but considering both complete and incomplete selective sweeps (selected allele frequencies ranged from 0.2 to 1). The age of selection *t* were randomly drawn from uniform distribution, t ∼ U(0, 100) kya. The distribution of the selection coefficient *s* is an equal mix between a Gamma distribution (60% of *s*≤0.01) and a L-shape distribution (90% of *s*≤0.01). (**A**) ABC estimates (*y*-axis) performed using 2×10^5^ ABC simulations using *X* (*x*-axis) randomly drawn in flat prior distributions, X ∼ U(0, 300). See also the legend of Fig. 2B. (**B**) Simulated ORs obtained for fixed values of *X*, X = 100, 150], in Europe (blue) and Asia (green). The empirical ORs are indicated in red (solid dots for the populations CEU and CHB). See also the legend of Fig. 1B. Note that the rRMSEs (Online Methods) shown in **A** are higher than assuming incomplete sweeps only (Fig. 2B) because the rate of increase of the OR with *X* is lower in panel **B** than assuming incomplete sweeps only (Fig. 1B). (**C**) Posterior distributions of *X* in each European and Asian population (CEU and CHB are indicated in bold). The estimations were performed using 2×10^5^ ABC simulations used in **A**. The point estimates and CIs obtained in each population including the African populations, are given in Supplementary Table S2.

### A trend toward more sweeps among non-African populations

As expected given the higher derived allele frequencies (DAFs) observed outside Africa (Coop, Pickrell, Novembre, et al. 2009) we found a non-significant trend toward younger selection of higher intensity in non-Africans (overlapping CIs of the *S* and *T* estimates, Fig. 3E-F, Supplementary Table S2). For example, our *S* estimates (obtained assuming incomplete sweeps only) were found higher in CEU than in YRI, i.e., 0.015 [CI:0.01-0.03] *vs* 0.09 [CI:0-0.02], in agreement with previous estimations (0.034 *vs* 0.022 in the same populations (Hawks, et al. 2007)). Our estimates are significantly lower likely because the selection models assumed differ (dominant model *vs* additivity in the present study). Moreover, our method may underestimate *S* when selection is strong as acknowledged above (Supplementary Fig. S3). We also found a trend toward more sweeps among non-African populations, particularly in Asia (Fig. 3A-C). For example, we estimated 68 [CI:46-91] and 165 [CI:119-211] sweeps in average in Africa and Asia respectively when including complete sweeps in our model (Supplementary Table S2). As the non-African estimates of *X_2_* and *X_3_* were consistently found higher than in Africa (Fig. 3D and Supplementary Figs. S18-19), our results also suggest more soft sweeps with low and intermediate initial frequencies (0.01≤*p_start_*<0.2, *X*_2_+*X*_3_) in Eurasian populations. Our estimations of higher numbers of sweeps in Eurasians populations are in agreement with greater numbers of sweeps detected outside Africa (Granka, et al. 2012; Pybus, et al. 2015; Schrider and Kern 2016).

### Selection signals in 1000G populations tend to be continent-specific

The method used here relies on genome-wide enrichment of candidate SNPs of selection in extended genic regions. We thus determined the genomic regions which contribute to this genome-wide enrichment, i.e., those enriched in candidate SNPs for the same neutrality statistics as used in our ABC estimations (Online Methods). We found that the most enriched genomic regions contain multiple iconic examples of positive selection, e.g., the *LCT* region in northern Europeans (Bersaglieri, et al. 2004), *EDAR* in all Asian populations and *TLR5* in Africa (Grossman, et al. 2013), as well as various other examples including genes associated with lighter skin pigmentation (Supplementary Text, see the most enriched genomic regions in Supplementary Table S3). This indicates that the information underlying our estimations corresponds to genomic signals in line with natural selection. We next investigated the overlap of these enriched genomic regions across populations (Online Methods) to roughly estimate the number of sweeps in humans as a whole. We found virtually no overlap between populations from different continents (Supplementary Fig. S20), with some exceptions that are indicated in Supplementary Fig. S21A and Supplementary Table S3. In contrast, the most enriched regions tend to be shared across populations from the same continent (Supplementary Fig. S20, Supplementary Table S3, an example given in Supplementary Fig. S21B). These results indicate that the total number of sweeps in humans should be close to the summation of the mean *X* computed per continent; i.e., 221 [CI:122-329] (Supplementary Table S2), a first order approximation neglecting some selection signals shared across continents and considering sweeps are highly shared within continents.

## Discussion

Our study revealed very limited numbers of selective sweeps in humans regardless the selection model considered (i.e., considering or not complete sweeps). We did not use McDonald–Kreitman-based methods since such methods aim to infer adaptation rate since divergence with other primate species (Uricchio, et al. 2019), and are potentially weakly sensitive to recent selective sweeps within species (Messer and Petrov 2013). Instead, we implemented an *ad hoc* ABC method to formally quantify numbers of selective sweeps over shorter time spans (∼100,000ya). This method does not scan genomic regions separately and thus cannot provide lists of genomics region under selection (normal genome-wide scans can be conducted in parallel, as we did in this study). It nevertheless helps to alleviate the load of false positives described in introduction (weakly or moderately stringent detection thresholds used to capture some sweeps of weak intensity provide large numbers of sweeps mainly explained by false discoveries while highly stringent thresholds used to discard false positives provide low numbers of detected sweeps poorly representative of the extent of selection operating in populations). Here, using selection signals at 1% threshold we showed that convenient statistics such as the OR for selection can provide suitable information to quantify real numbers of selective sweeps in populations. Note that a similar ABC method can be implemented using more than six ORs when background selection is explicitly simulated in the ABC model, including ORs computed for *F*_ST_, AFS-based statistics and others (the expected gain in accuracy remains to be formally assessed).

Our estimations are not perfect and can still be improved. For example, we may have missed some selective sweeps hidden in regions not yet classified as functional (selection signals found in empirical ENVs downward bias the *X* estimations by decreasing the empirical ORs). However, 70% of the genome was considered as potentially influenced by selection and all significant signals of positive selection found in intergenic regions have been accounted for (e.g., Supplementary Fig. S18), suggesting that we captured a large fraction of the existing selective sweeps. Simulating whole chromosomes is still barely tractable in term of computation times. We thus concatenated shorter simulated regions together, trimming their edges and avoiding computation of any neutrality statistics across the junctions. This insures that haplotype-based statistics were computed over linked SNPs simulated according to human recombination rates. The demographic model assumed incorporates major components of human demography (e.g., expansion, bottleneck and isolation-with-migration) but some other aspects are ignored, such as the admixture with ancient hominids. However, we found low differences in simulated ORs obtained with contrasting demographic histories, we found similar estimations of *X* when replacing an expansion with a bottleneck, and only slight overestimations of *X* when the bottleneck assumed is four times stronger than in reality.

Although we do not wish to claim that we have found the exact numbers of selective sweeps (as if the demography was exactly known), our results indicate that the limited numbers of sweeps estimated herein are not due to our demographic assumptions. Note that discrepancies between point estimates across populations of the same continents must be interpreted with cautious in term of selection (point estimates with highly overlapping CIs should be considered as indistinguishable). They are likely explained by local demographic specificities that are not properly accounted by the model (e.g., *X* was sometime found lower in FIN than in CEU potentially because of a stronger bottleneck in the Finnish population (Bulik-Sullivan, et al. 2015)).

A main result of our study is the very limited numbers of selective sweeps inferred over the last ∼100,000 years, even when modelling both complete and incomplete sweeps. It is worth mentioning that our method can estimate *X* despite being based on neutrality statistics that cannot even be computed when the selected alleles are fixed, e.g., iHS (even lower, ORs are informative because the relationship with *X* is maintained). This illustrates the importance of defining as PSVs the SNPs nearby selection targets because our method uses information at linked sites through the enrichment of candidate SNPs around the fixed selected alleles. Assuming similar sweep completion times (ranging from 0 to 3,000 generations) and similar initial frequencies of selected alleles (ranging from 1/2*N* to 0.2), we estimated a much lower number of selective sweeps in Europe and Asia (115 [CI:68-160] and 165 [CI:119-211] respectively) than previously detected in CEU (∼1,013 sweeps) and JPT (∼1,059 sweeps) (Schrider and Kern 2017). Beside our estimations, ∼1,000 sweeps should result in simulated ORs too large to be compatible with the empirical ORs observed (Fig. 1B, Fig. 4B), suggesting that the majority of these predicted soft sweeps can be explained as mis-classified neutral regions, as pointed by (Harris, et al. 2018). Nevertheless, even though we voluntary considered the same time span as used by Schrider and Kern in their training simulated datasets we may have missed some older selection signals that could be captured by their machine learning algorithm (our CIs should be enlarged when considering older selection ages in our ABC model). In addition, we also estimated, except in Africa, lower numbers of sweeps than previously detected based on another machine learning algorithm (355 and 424 detected sweeps in CEU and CHB respectively (Pybus, et al. 2015)). Apart from the sensitivity to false positives of this method based on detected/classified sites, such discrepancies are difficult to interpret because the machine learning algorithm has been trained using hard sweeps only, ignoring the soft sweep model.

More generally, we showed that our method can easily be extended to incorporate complete sweeps. In doing so, we estimated ∼twice more selective sweeps in Eurasia than assuming complete sweeps only, e.g., 165 vs 88 sweeps in Asia (Supplementary Table S2). Our results thus suggest limited numbers of complete sweeps over the last ∼100,000 years (e.g., at most 165 complete sweeps in Asia) and confirm that recent human adaptation has also been driven by a significant fraction of incomplete sweeps, some of them potentially ongoing (Wilde, et al. 2014; Field, et al. 2016). (Nearly all iconic sweeps are associated with a putative advantageous mutations still segregating in current generations (Vitti, et al. 2013; Jeong and Di Rienzo 2014; Fan, et al. 2016)). A similar ABC approach can be further designed to formally estimate the numbers of complete and incomplete sweeps specifically.

We also estimated a majority of sweeps with initial frequencies greater than 1% (0.01≤*p_start_*<0.2, *X*_2_+*X*_3_), the rest being sweeps on very rare standing variants (1/2*N≤p_start_*<0.01, *X*_1_). Given the known difficulty to distinguish between hard and soft sweeps even for initial frequencies ranged to 0.2 (see some high miss-classification rates in (Schrider and Kern 2017; Harris, et al. 2018)), we did not provide separate estimates of the numbers of hard and soft sweeps on very rare standing variants. The numbers of hard sweeps remain unknown and may thus range from 0 to the estimated value of *X*_1_. However, our results (*X*_1_<*X*_2_+*X*_3_) confirm soft sweeps as main drivers of human adaptation relative to hard sweeps (Schrider and Kern 2017) (some examples have already been described in humans (Peter, et al. 2012) and also in drosophila (Garud, et al. 2015), the latter have been discussed in (Harris, et al. 2018)). Nevertheless, we only observed a moderate excess of sweeps on standing variants with frequency higher than 1% (*X*_2_+*X*_3_≈2/3*X*) because the proportion of mutations with frequency greater than 1% drastically drops, as predicted by theory and consistent with what is observed in extant populations (a mutation becomes advantageous irrespective to its frequency). It is worth mentioning that we also found greater numbers of such soft sweeps outside Africa (see the *X*_2_ and *X*_3_ estimated, Fig. 3D and Supplementary Fig. S19A). It is not necessary to assume biological features or changes in selective regimes in Eurasia since the proportion of such sweeps also depends on the demography of the populations considered. Indeed, the loss (by genetic drift) of selected alleles at very low frequency during the first generations of selection may be exacerbated in bottlenecked populations relative to expanding populations. This may causes diminished numbers of sweeps on very rare variants in Eurasian populations relative to Africa, and consequently, a mechanical excess of sweeps on the other standing variants (with frequency higher than 1%) outside Africa.

We would-like to emphasize that the scenario of positive selection on standing variation investigated here is not related to polygenic adaptation (Pritchard and Di Rienzo 2010; Pritchard, et al. 2010). Because biological processes likely differ (very few advantageous alleles with large phenotypic effects *vs* many advantageous alleles with small phenotypic effects), soft sweeps drive to fixation while polygenic adaptation refers to moderate changes of selected allele frequencies only (Pritchard, et al. 2010). Hence, the neutrality statistics used in this study, which are poorly sensitive to small changes in allele frequencies, do not capture the pervasive effects of polygenic adaptation in humans (Field, et al. 2016). Instead, the number of selective sweeps we report should be interpreted as the minimum number of sweeps having occurred in a population. For example, given the important reproductive function of the *SPAG4* gene (Kracklauer, et al. 2010), the species-wide selection signal found near this gene (Supplementary Fig. S21A) is consistent with a single selection hit in each continent independently. However, other functional variants found near the *SPAG4* region suggest a more complex scenario in Asia, involving several independent selection hits in the Asian lineage specifically, e.g., regulatory variants of *GDF5* have also been found under selection in Asia (Capellini, et al. 2017).

Finally, the higher numbers of sweeps we estimated in non-African populations support models in which more adaptation is expected when populations colonize new environments. In an already colonized environment, populations also need to adapt to environmental pressures that can change over time. The continent-specific signal of selection shared among all European populations, encompasses eQTLs downregulating the expression of the *PLEK2* gene (Supplementary Fig. S21B) in skin cells exposed to the sun (GTEx database, (Ardlie, et al. 2015)). The cold paleoclimate (last ice age, from ∼110 to ∼10kya (Cooper, et al. 2015)) favored both light skin pigmentation and increased sensitivity to UV-induced melanoma in Europe (Lopez, et al. 2014; Key, et al. 2016) while low expressions of *PLEK2* may increase the survival probability in melanoma patients (Supplementary Text). The alleles downregulating *PLEK2* could have been favored by selection during the late Pleistocene climatic warming (∼20 to ∼10kya), as supported by selection signal of ∼1-1.2Mb suggesting a recent onset of selection that may coincide with the increase of regional temperatures and solar luminosity in the northern hemisphere (Luo, et al. 2011) (Supplementary Text).

Our study provides independent lines of evidence that selective sweeps, despite their small numbers with respect to other modes of adaption such as negative selection or polygenic adaptation, enabled human populations to adapt to environmental pressures. The method proposed can be applied in other species (e.g., Drosophila (Garud, et al. 2015), mouse (Ihle, et al. 2006), domesticated animal breeds (Stella, et al. 2010; Roux, et al. 2015), domestic dogs (Freedman, et al. 2016) and horses (Librado, et al. 2015)) and extended to other modes of selection (e.g., adaptive introgression from ancient hominids or short-term balancing selection), allowing for broader investigations of the impact of extreme environments, domestication or mode of reproduction on recent adaptive evolution of species.

## Materials and Methods

### 1000 Genomes populations analyzed

Analyses were performed on the 1000 Genomes Project phase 3 data, focusing on African, European and Asian populations. We analyzed 1,511 individuals from five African, five European and five East-Asian populations (85 to 113 individuals per population, Supplementary Table S1), excluding populations with diverse continental or admixed ancestry, for which little demographic history information was available. We downloaded phased data obtained with SHAPEIT2 (Delaneau, et al. 2012), ancestral/derived states and VEP annotations from the 1000G Project website.

### Simulating WGSs

We carried out computer simulations of WGSs with various *X* values, ranging from 0 (neutrality) to >200 per population. The simulated WGSs mimicked the 1000G data, with millions of SNPs spread over the genomes of ∼100 unrelated individuals per population. As we focused on the adaptive history of African, European and Asian populations in this study, we systematically simulated triplets of populations according to inferred models of African, European and Asian demography: the population with *X* loci under positive selection and two other neutral populations (*X*=0), used as a reference for computations of interpopulation neutrality statistics. Specifically, we used a three-populations isolation-with-migration model (the parameters used can be found in (Grossman, et al. 2013)). This model incorporates an ancient African expansion, an Out-of-Africa exodus ∼100 kya (28 years per generation (Fenner 2005; Moorjani, et al. 2016)) followed by a bottleneck and a split of Eurasians into European and Asian populations ∼58kya. This model also reproduces different migration rates between continents, with a probability of the order of 10^-5^ per haploid genome per generation. One key feature is the presence of two population bottlenecks in non-African populations, the second bottleneck being stronger in the Asian population (Pickrell, et al. 2009b).

We used SLiM (Haller and Messer 2017) to simulate 5Mb regions sequenced in 100 unrelated individuals per population, using human recombination rates sampled from the HapMap recombination maps (Frazer, et al. 2007). We simulated 10^4^ neutrally evolving DNA regions and 2×10^3^ selected DNA regions. For each triplet of populations, we simulated selection in the population of interest (African, Asian or European) by inserting (in the middle of the region) an advantageous mutation at generation *t*, with a frequency, *p_start_*, ranging from 1/2*N* to 0.2 as previously assumed (Schrider and Kern 2017). The *p_start_* was randomly drawn from the allele frequency spectrum at the generation *t*. The intensities and ages of selection were randomly drawn from specified distributions (see main text). It is worth mentioning that we simulated long DNA regions to avoid premature truncation because selection signals can extent over mega bases for selection events, particularly recent and/or strong events, i.e., ∼2Mb in the LCT region (various estimates of s for rs4988135 ranged from 0.025 to 0.069 (Tishkoff, et al. 2007; Peter, et al. 2012; Chen, et al. 2015)). Because SLiM is a forward-in-time simulator, the computation times, which depend on both the effective population size *N* and the *t* generations simulated, are large for the model investigated. We therefore optimized computational times by rescaling effective population sizes and times according to *N/λ* and *t/λ* with *λ* = 10 (Hoggart, et al. 2007). We used rescaled mutation and recombination rates, *λµ* and *λr*. Similarly, because we divided the number of generations by *λ*, the selection parameter s must be multiplied by the same factor (Hoggart, et al. 2007; Haller and Messer 2017). Finally, WGSs were obtained by concatenating randomly drawn 5Mb regions, some of which were considered to be ENVs, the rest being PSVs (see below). We built WGSs by restricting the *X* simulated sweeps to PSV regions, because ENV genomic regions are, by definition, neutral. The ABC simulations used for estimations are simulated sets of such WGSs (10^5^ simulated WGSs, the prior distributions used for *X*, *S* and *T* are specified below).

### PSVs in simulated WGSs

PSVs (Supplementary Fig. S1A) are all sites potentially influenced by selection either directly (all potential targets of selection, i.e. all variants altering phenotypes in real data) or indirectly (all neutral SNPs in the vicinity of a potential target of selection). The probability *ξ_i_* that the *i^th^* SNP is influenced by selection starts from 0 (neutral SNP far from the potential targets of selection) and increase to 1 when approaching a potential target of selection. In this analysis, we discretized this probability by considering an indicator variable *I*_i_, assigned to SNP *i* and equal to 1 (PSV) or 0 (ENV):

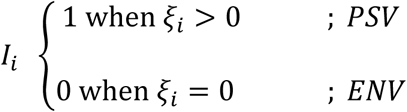

In simulated WGSs, the PSVs are randomly simulated as tracts of SNPs with *I*_i_=1 randomly spread over the genomes. We reproduced the same proportion of PSVs as observed in 1000G populations, with numbers of simulated SNPs per population matching those observed in 1000G populations. In our simulations the recombination rates are randomly sampled in human recombination maps and are thus similar across PSVs and ENVs. However, in real data the recombination rate is lower in genes than in intergenic regions. Initially lower in genes it suddenly increases in flanking regions and progressively decreases with distance from the nearest gene to become lower in remote intergenic regions (Myers, et al. 2005; Coop, et al. 2008; Kong, et al. 2010), making our simulations conservative with respect to the rates of candidate SNPs in ENVs far from genes. We however also tested higher recombination rates in ENVs than in PSVs (Kong, et al. 2010) and re-estimated the parameters accordingly (supplementary Fig. S18).

### Neutrality statistics and candidate SNPs in simulated WGSs

For each simulated WGS, we computed several widely used neutrality statistics expected to have extreme values for SNPs targeted by selection or located close to such SNPs. We used haplotype-based neutrality statistics, which compare the haplotypes carrying the derived and ancestral alleles, iHS, DIND, ΔiHH, and the derived alleles between populations, XP-EHH. We also used the Fay and Wu’s *H*, which detect deviations from the neutral allele frequency spectrum (AFS) in short genomic regions. We used a sliding-windows approach (100 kb windows centered on each SNP (Fagny, et al. 2014)) for these computations. The sliding-windows began and ended 50kb from the edges of the 5Mb simulated regions, to prevent truncation in the 100 kb sliding windows (a similar approach was applied to the 1000G chromosomes). As iHS, DIND, ΔiHH and XP-EHH are sensitive to the inferred ancestral/derived state, we computed these statistics only when the derived state was determined unambiguously (Fagny, et al. 2014) (i.e. more than 95% of SNPs). We then normalized these statistics by DAF bin (Voight, et al. 2006; Fagny, et al. 2014) (mutations grouped by DAF bin, from 0 to 1, in increments of 0.025). We minimized the false-positive discovery by excluding SNPs with a DAF below 0.2, as the power to detect positive selection has been shown to be limited at such low frequencies (Voight, et al. 2006; Fagny, et al. 2014). The method used are implemented in selink, a software to detect selection using whole-genome datasets (https://github.com/h-e-g/selink). Finally, for each neutrality statistic, we defined candidate SNPs of selection as the 1% of SNPs with the most extreme values over 10^4^ neutral simulations. For iHS, DIND, ΔiHH and XP-EHHs, we considered extreme values indicative for selection targeting the derived alleles, e.g, large negative values of iHS indicate unusually long haplotypes carrying the derived allele.

We choose these neutrality statistics because the widespread effects of background selection may be confounded with positive selection as mentioned in the main text. For example, the Tajima’s D is sensitive to BGS (Zeng, et al. 2006) and the differences in allele frequencies between-population are expected to be exacerbated in regions affected by BGS, a pattern that can be confounded with positive selection (Coop, Pickrell, Novembre, et al. 2009; Pickrell, et al. 2009b; Pritchard, et al. 2010). We excluded *F*_ST_, Tajima’s D and others similarly affected by BGS. We only retained Fay and Wu’s H and the haplotype-based statistics previously found to be insensitive to BGS (Zeng, et al. 2006; Fagny, et al. 2014), e.g., haplotype-based statistics compare haplotypes carrying the derived allele with those carrying the ancestral alleles as internal controls that should be affected by BGS to a similar extent. We used this approach (excluding statistics sensitive to BGS) rather than simulating widespread BGS, because we aimed to avoid bias in our ABC estimations due to the imperfectly known patterns of BGS on the human genome.

### Odds ratio for selection used as ABC summary statistics

The summary statistics used in the present study were the odds ratio for selection (Kudaravalli, et al. 2009; Fagny, et al. 2014), computed for the population of interest (with *X* simulated or estimated), for each WGS (simulated or 1000G) and each neutrality statistic separately. Each OR assesses the enrichment of candidate SNPs in PSVs, as follows:

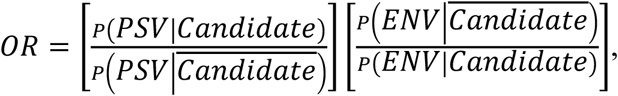

with Candidate and 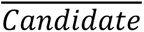 being candidate and non-candidate SNPs, respectively (see above). The OR is greater than 1 if positive selection has preferentially occurred in functional regions, because such selection tends to increase the number of candidate SNPs across the selective sweep regions (Voight, et al. 2006).

### ABC prior distributions, acceptance rules and accuracy validation

We randomly draw *X* from a uniform prior distribution *X* ∼ *U*(0, > 200), where *X*=0 is a neutral WGS. Priors for mean intensity and age, *S* and *T*, were derived from the priors for *s* and *t* used to simulate the *X* sweeps (the priors used are indicated in main text). We used the ‘abc’ R package and the standard ABC method (Beaumont, et al. 2002), in which posteriors are constructed from simulated parameters accepted and adjusted by local linear regression (method=“Loclinear” in the ‘abc’ package). The accepted simulated parameters are those which provide the best fit with empirical data (the ‘abc’ parameter ‘tol’ was set to be equal to 0.005). We used the mean of the posterior distribution as point estimate and provided the 95% credible interval boundaries (computed from the posterior distribution).

To test the accuracy of our estimations, we compared the estimated and simulated parameter values, *θ̂*_i_ and *θ_i_* respectively, using classical accuracy indices: the relative error *rE* (i.e. difference between estimated and true values, expressed as a proportion of the true value, rE = (*θ̂_i_* – *θ_i_*)/*θ_i_*, *i* = 1, …,*J*), the relative root of the mean square error, *r*RMSE (i.e. the root of the MSE expressed as a proportion of the true value), and the proportion of true values within the 90% credible interval of estimates, 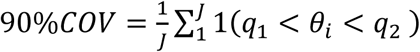 where 1(*C*) is the indicative function (equal to 1 when *C* is true, 0 otherwise) and *q*_1_ and *q*_2,_ the corresponding percentiles of the posterior distributions.

### Defining PSVs in the 1000G data

The definition of the PSVs in the 1000G populations is based on the VEP (Ensembl Variant Effect Predictor, Supplementary Fig. S8) annotations provided by the 1000G website. We included in PSVs the potential targets of selection as exhaustively as possible. We considered as PSVs missense variants, stop-gained mutations and all the regulatory variants, including SNPs located in the regions 5’ and 3’ to genes (regions expected to be enriched in eQTLs) together with the transcription factor binding sites located farther away from genes. As in simulations, we included all SNPs located in the vicinity of these potential direct targets of selection to take into account the hitchhiking effects of selection at nearby linked loci. For example, in the case of the high-altitude adaptation in humans, two of the largest Tibetan-Han frequency differences found in EPAS1 were annotated as intronic (Yi, et al. 2010). Hence, all synonymous, intronic, 5’ and 3’UTR SNPs (i.e., ∼50% of annotated SNPs) are included in PSVs. We also included upstream/downstream SNPs in PSVs to prevent early truncations of selective sweep signals. With such filters, ∼70% of 1000G SNPs were considered to be PSVs.

Finally, some particular cases are hardly tractable based on VEP annotation only. These include the selected sites that are unknown functional variants annotated as intergenic, small regulatory regions located far from other PSVs and regulatory variants located in edges of PSVs tracts. In these situations, selection signals may be found in ENV and therefore can bias downward our estimations (selection signals found in ENV cause lower empirical ORs). To minimize such estimation biases, we annotated as PSVs all the SNPs with a genome-wide significant enrichment in candidate SNPs (this enrichment is computed using 100 kb centered on every SNP, see below). High enrichment are indicative of selection, since, as stated above, positive selection produces clusters of candidate SNPs (Voight, et al. 2006). This step has only a marginal effect on the total number of SNPs classified as PSVs (∼0.01% of SNPs reallocated), but it can inflate neutral ORs (e.g., OR computed for iHS from purely neutral simulations, *X*=0, is equal to 1.2 rather than 1). We therefore reproduced this step in the simulations used to obtain ABC estimations for 1000G populations, i.e., the simulated SNPs annotated as PSVs included all simulated SNPs with a significant enrichment in candidate SNPs. We also performed a round of estimation after removing from the analysis all ENVs with a significant enrichment in candidate SNPs of selection (supplementary Fig. S18).

### Odds ratio for selection in the 1000G populations

When analyzing the 1000G populations, we computed the ORs using a logistic regression to control for several covariates (Kudaravalli, et al. 2009), including genomic variation in sequencing quality, recombination and mutation rates. For each neutrality statistic and chromosome, we set

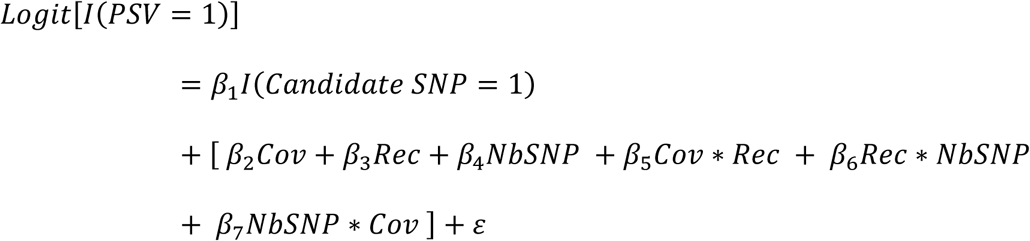

with I(*PSV* = 1), the indicator function, equal to 1 if the SNP is a PSV, or 0 otherwise, I(*Candidate SNP* = 1) being an indicator function equal to 1 if the SNP is a candidate SNP of selection or 0 otherwise (candidate SNPs are the 1% of SNPs with the most extreme values genome-wide). Rec is the mean recombination rate in cM/bp (obtained from HapMap recombination maps), Cov is the mean coverage, and NbSNP is the number of SNPs computed with 100 kb sliding windows. The OR was estimated with exp(*β*_1_) and averaged over chromosomes (Kudaravalli, et al. 2009). We also performed a round of estimation using empirical ORs computed merging all chromosomes together (supplementary Fig. S18).

### Assessing the genomic regions enriched in candidate SNPs of selection

For each SNP and each of the six neutrality statistics used in the ABC estimations (see main text), we computed the proportion of candidate SNPs in a 100 kb window around the SNP being considered. We next determined the empirical *P*-values (*P*) for these proportions and combined them into a single combined selection score (denoted by CSS) using a Fisher score, 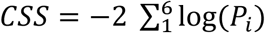 (Deschamps, et al. 2016). The rationale behind such a composite approach (Grossman, et al. 2010; Grossman, et al. 2013; Deschamps, et al. 2016) is that neutrality statistics are more strongly correlated for positively selected variants than for neutral sites (Grossman, et al. 2013). Consequently, false positives may harbor extreme values for a few neutrality statistics only, whereas SNPs genuinely selected (or nearby SNPs) should harbor extreme values for several statistics together, a feature captured by the combined score. Finally, the genomic regions enriched in candidate SNPs were defined as consecutive SNPs with genome-wide significant CSS values (*P*<0.01). In these enriched regions, the strength of selection signal has been related to the magnitude of the enrichment in candidate SNPs through the maximum CSS value found in the region. Indeed, high combined selection score values are expected for SNPs targeted by young and strong selection and for the others SNPs nearby, e.g., SNPs nearby rs4988235 (*LCT* region) were found among the highest CSS values found genome-wide (Supplementary Table S3).

### Assessing the overlap of selection signals between the 1000G populations

For each region enriched in candidate SNPs of selection, we assessed the sharing of selection signals between populations by mean of an overlap score, calculated as the number of populations for which the same enriched region was identified. For each enriched region, overlap scores were calculated both within and between continents (upper limits of 5 for within-content and 10 for between-continent scores, because there were five populations from each of three continents). Thus, continent-specific enriched regions would have within- and between-continent overlap scores of 5 and 0 respectively, whereas population-specific enriched regions would have within- and between-continent overlap scores of 1 and 0, respectively.

## URLs

1000 Genomes data: ftp://ftp.1000genomes.ebi.ac.uk/vol1/ftp/release/20130502/, ancestral state, ftp://ftp.ncbi.nih.gov/1000genomes/ftp/technical/working/20120316_phase1_integrated_release_version2/, VEP functional annotation, ftp://ftp.ncbi.nih.gov/1000genomes/ftp/technical/working/20120316_phase1_integrated_release_version2/. tools for Approximate Bayesian Computation (ABC), http://cran.r-project.org/web/packages/abc/index.html, GTEx (Genotype-Tissue Expression Project) http://gtexportal.org/home/, DAA (Digital Aging Atlas) http://ageing-map.org/, the human protein atlas http://www.proteinatlas.org, the American Cancer Society https://www.cancer.org/, NCBI gene database https://www.ncbi.nlm.nih.gov/gene/26499

## Supplemental Data

Supplemental data, including Figs. S1 to S21 and Tables S1 to S3 (Tables S2 and S3 are supplied as Excel files) can be found with this article online.

## Acknowledgments

This work was supported by the Institut Pasteur, the *Centre Nationale de la Recherche Scientifique* (CENV), and the *Agence Nationale de la Recherche* (ANR) grants: “DEMOCHIPS” ANR-12-BSV7-0012, “IEIHSEER “ ANR-14-CE14-0008-02 and “TBPATHGEN” ANR-14-CE14-0007-02. The laboratory of LQM has received funding from the French Government’s Investissement d’Avenir program, Laboratoire d’Excellence “Integrative Biology of Emerging Infectious Diseases” (grant no. ANR-10-LABX-62-IBEID), and from the European Research Council under the European Union’s Seventh Framework Programme (FP/2007–2013)/ERC Grant Agreement No. 281297.

## Author contributions

G.L. designed and performed computational analysis, analyzed and interpreted results, and wrote the paper. P.B. designed and optimized the selink code. E.P. and L.Q.M. advised on analysis and data interpretation and helped in writing the manuscript.

## Corresponding authors

Correspondence to: Guillaume Laval at

