## Supplemental Information for "A genome-wide Approximate Bayesian Computation approach suggests only limited numbers of soft sweeps in humans over the last 100,000 years"

**Figure S1. Possibly selected variants (PSVs)**

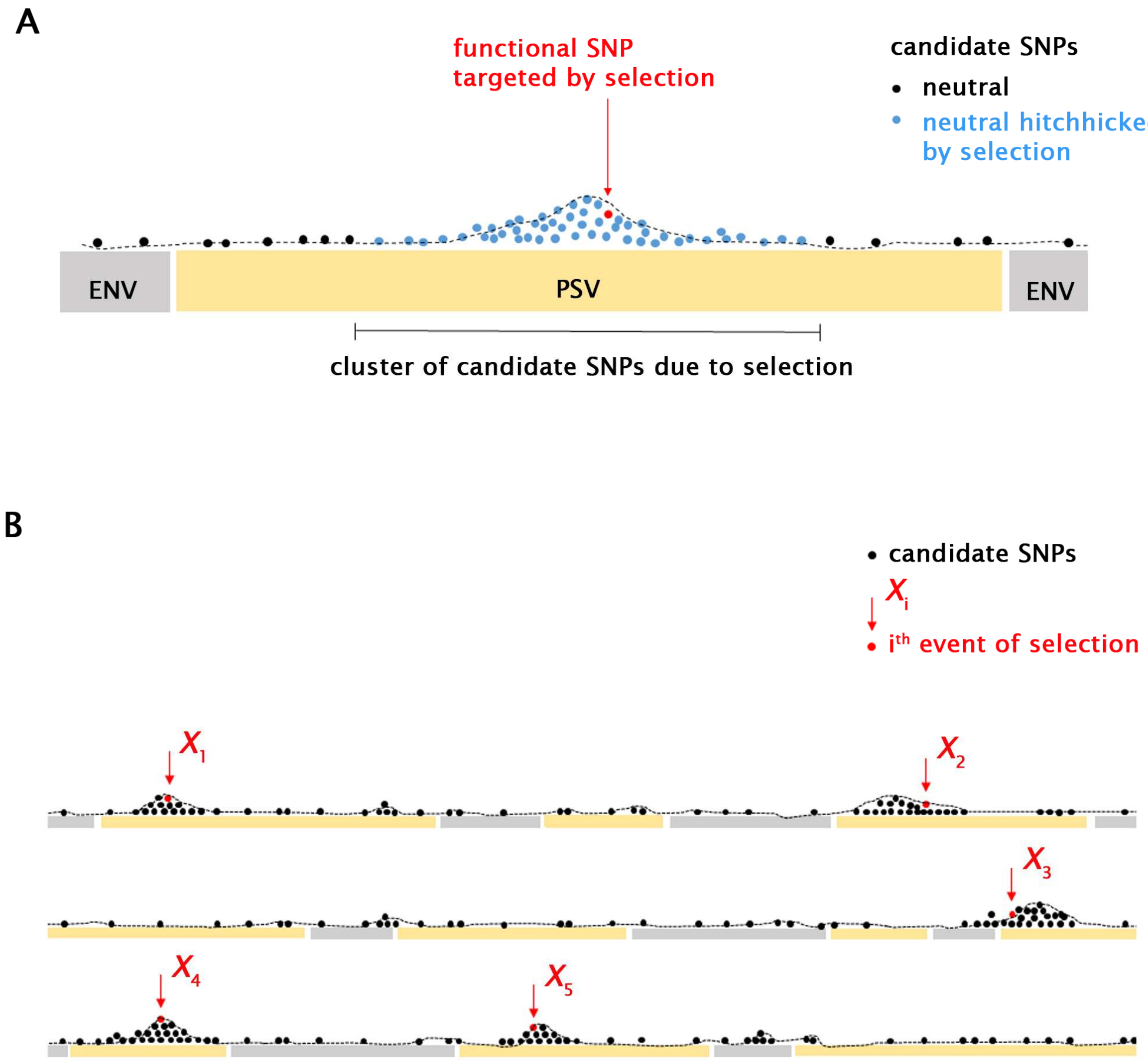


(**A**) PSVs (possibly selected variants) are all potential targets of selection and neutral nearby SNPs: every functional variant is potentially a direct target of selection (red dot, e.g., non-synonymous and regulatory variants close to genes) while among the neutral nearby variants (e.g., non-synonymous, intronic variants, intergenic variants, etc.) some of them are close enough to hitchhike during selection (blue dots) and others are not affected by selection (black dots). ENVs (evolutionary neutral variants) are unaffected by selection (e.g., intergenic SNPs at distance from the nearest gene and purged from any functional sites). (**B**) Example of $X$=5 selective sweeps (targeting the SNPs indicated in red) that can be simulated, highlighting the expected enrichment of candidate SNPs in PSVs.

**Figure S2. Simulated ORs for different categories of selective sweeps**

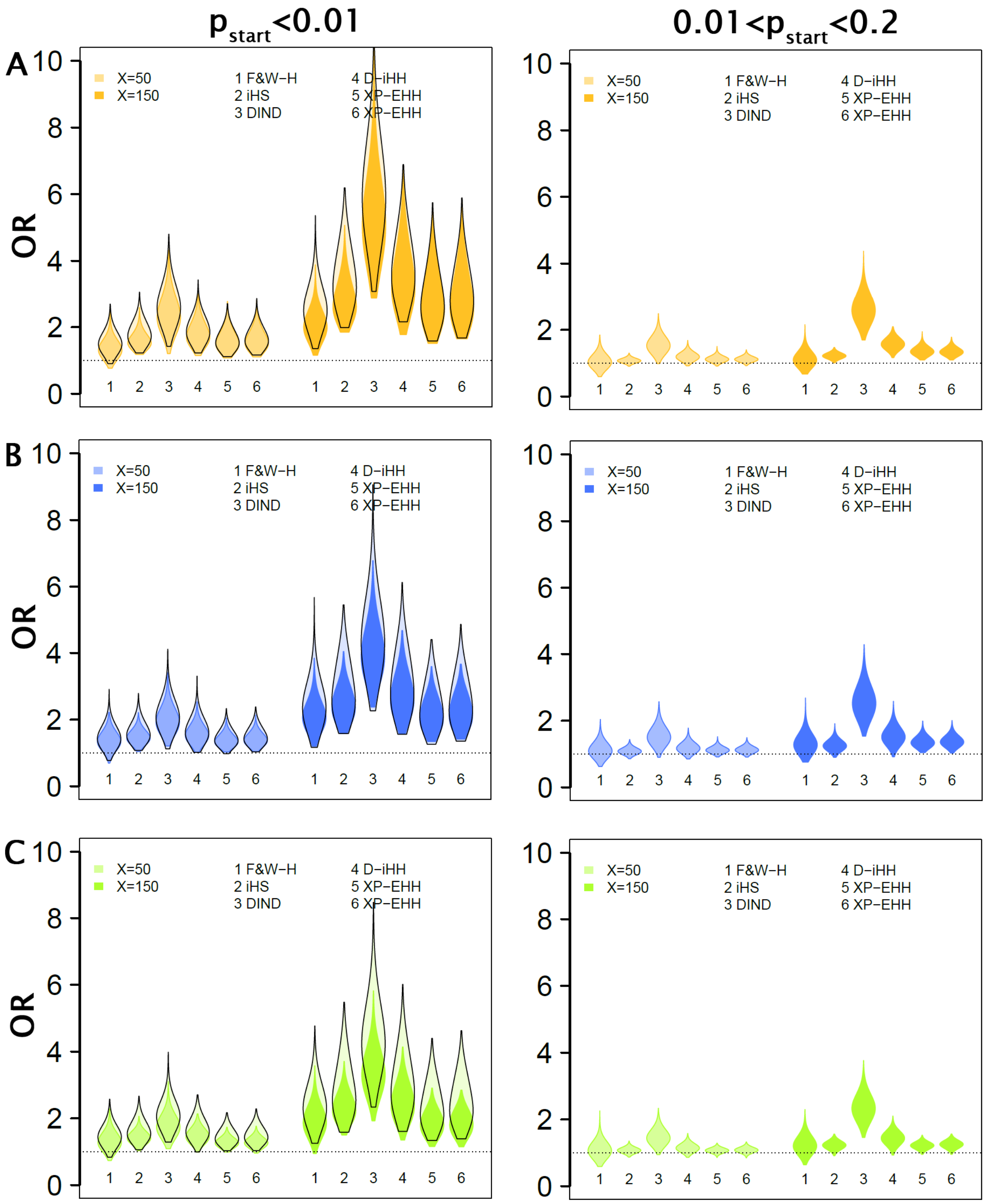


Relationship between the odds ratio for selection (OR) and the number of selective sweeps ($X$), in African (**A**), European (**B**) and Asian (**C**) simulated populations. The parameters underlying the simulations are described in Fig 1. (left panel) Selective sweeps simulated using frequency at the onset of selection ranged from $1/2N$ to 0.01. The distribution of ORs obtained when simulating hard sweeps *stricto senso* (frequency at the onset of selection equal to $1/2N$) are indicated for comparison (empty violin plots with contours in black). (right panel) Selective sweeps simulated using frequency at the onset of selection ranged from 0.01 to 0.2.

**Figure S3. Cross-validation of the estimation of *S* and *T***

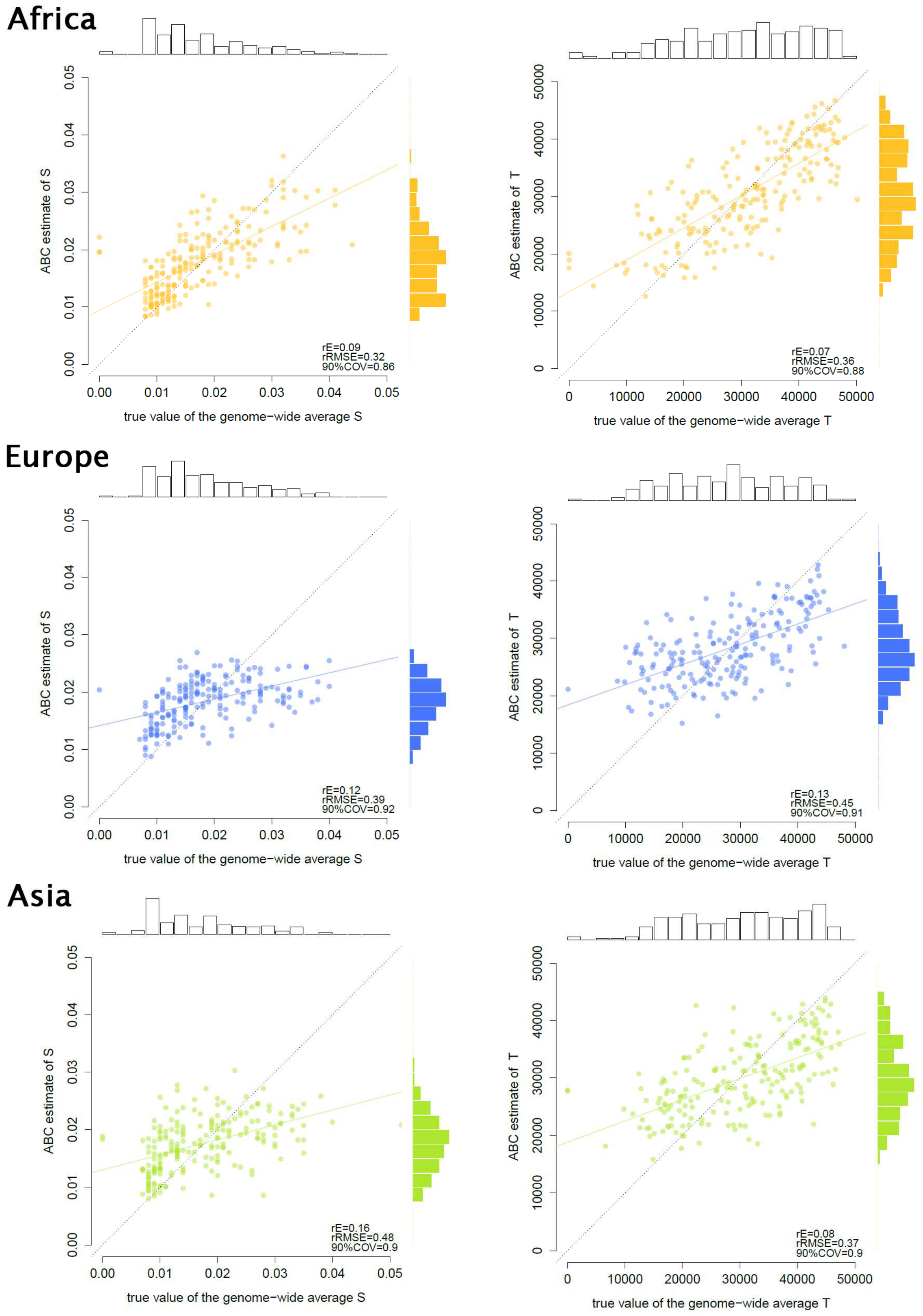


ABC estimations performed with the simulated WGSs used as empirical data shown in Fig. 2B. See the legend of Fig. 2. Prior distributions used for the intensity *s* and age of selection *t* of each sweep are shown in Fig. 1A.

**Figure S4. Cross-validation of the estimation of S and T**

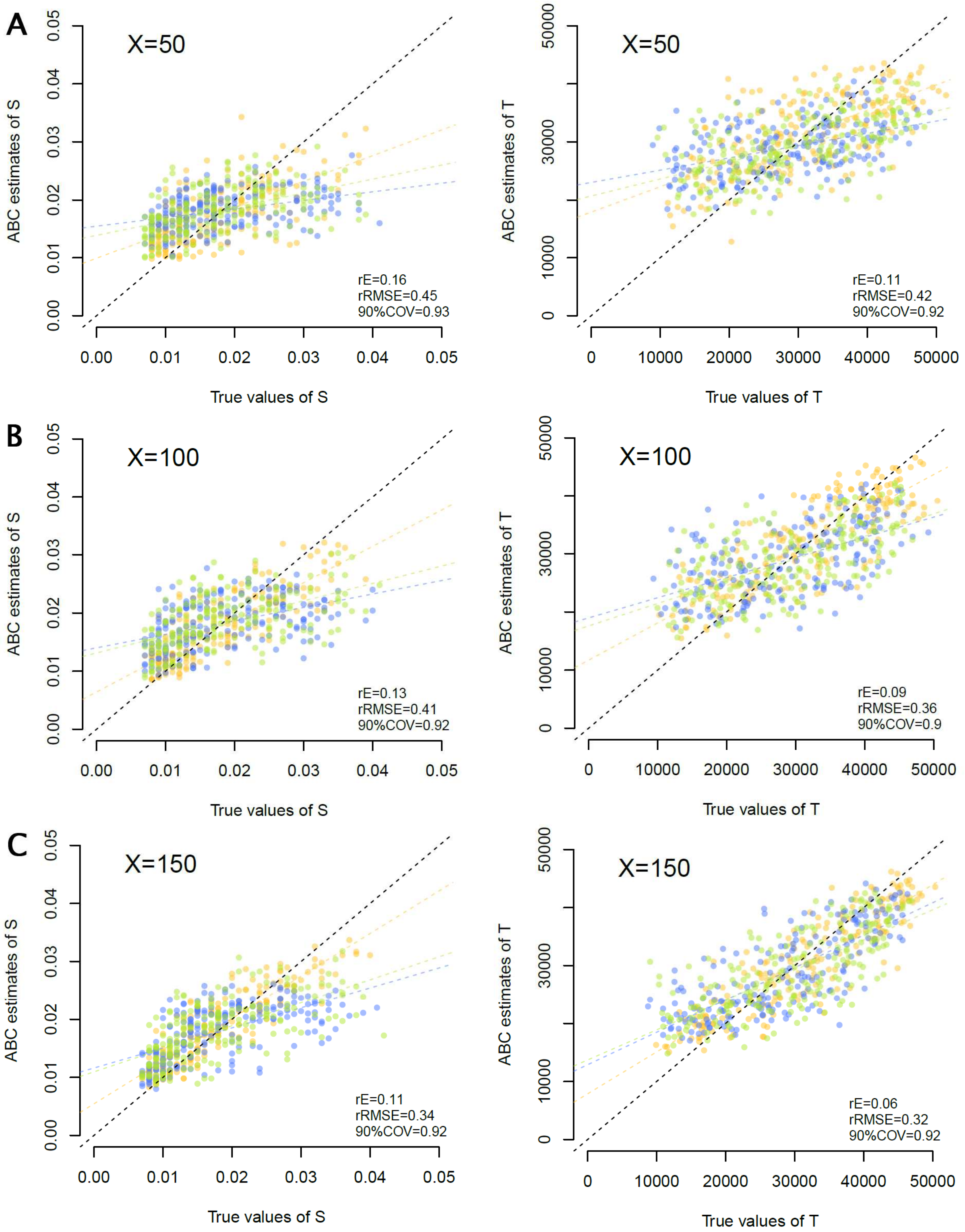


ABC estimations performed with the simulated WGSs used as empirical data shown in Fig 2A. Prior distributions used for the intensity *s* and age of selection *t* of each sweep are shown in Fig. 1A. African (yellow), European (blue) and Asian (green) simulated WGSs. The regression lines computed separately for each population show that the accuracy of the estimations are nearly identical in the various simulated populations, and, as expected, increase with the number of selective events.

**Figure S5. No correlation between *X* and *S* or *T* estimates**

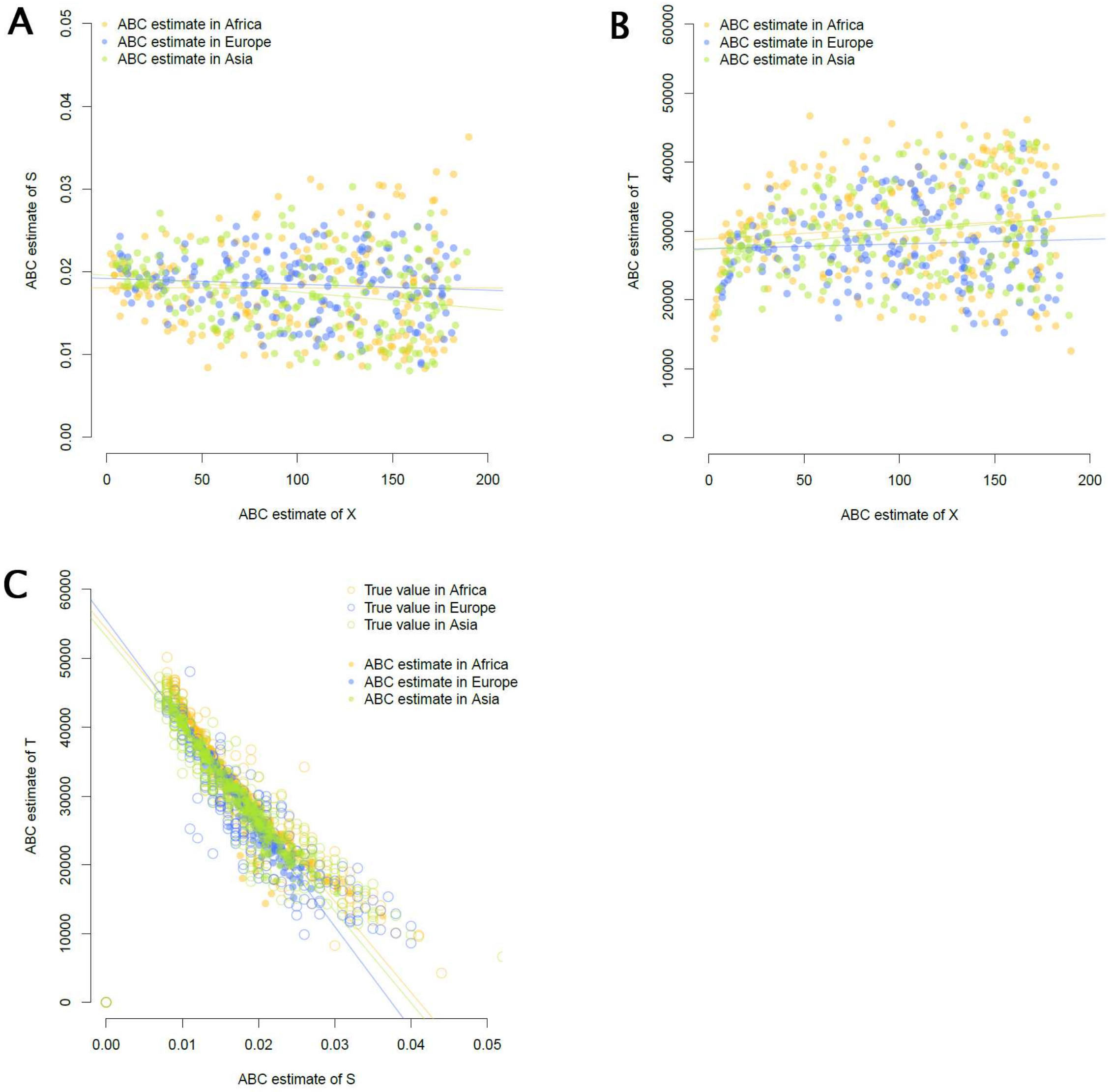


The ABC estimates are those shown in Fig. 2B for *X* and in Supplementary Fig. S3 for *S* and *T*. (**A-B**) No significant correlation between the estimations of *X* and the estimations of *S* or *T*. (**C**) Significant correlations between the estimations of *S* and *T* due to expected correlations in the model itself (true values in the model are indicated by empty circles). Regression lines in Africa, Europe and Asia are indicated in the corresponding colors.

**Figure S6. Credible intervals of the estimations of the *X*s.**

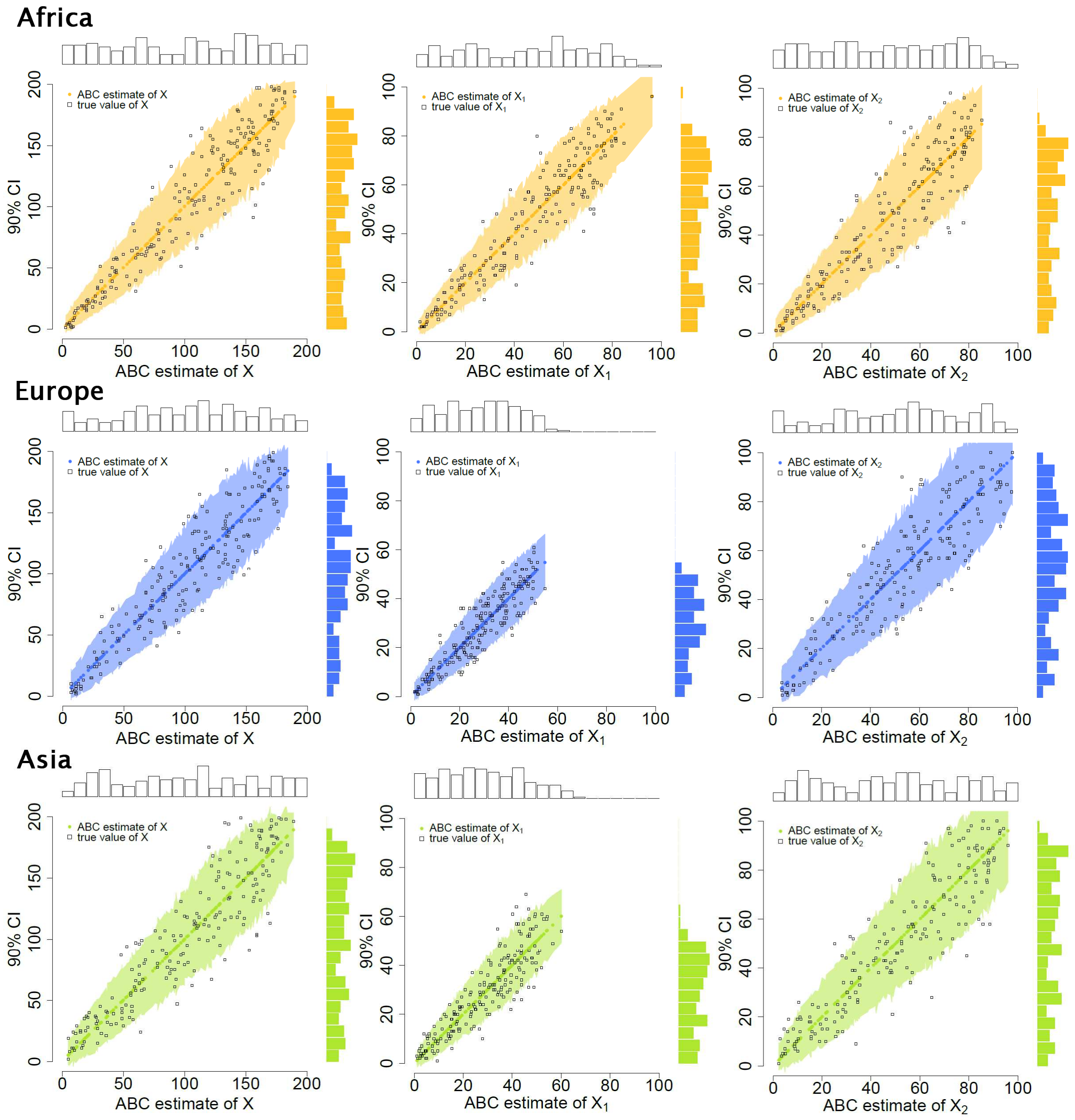


The ABC estimates are those shown in Fig. 2B. Prior distributions used for the intensity *s* and age of selection *t* of each sweep are shown in Fig. 1A. Range of 90% CIs (credible intervals, y-axis) plotted against the ABC estimations (*x*-axis). Black squares are the true values of the corresponding parameters. The external bar plots show the distributions of the true (top bars) and estimated (colored bars on the right) parameter values.

**Figure S7. Credible intervals of the estimation of *S* and *T*.**

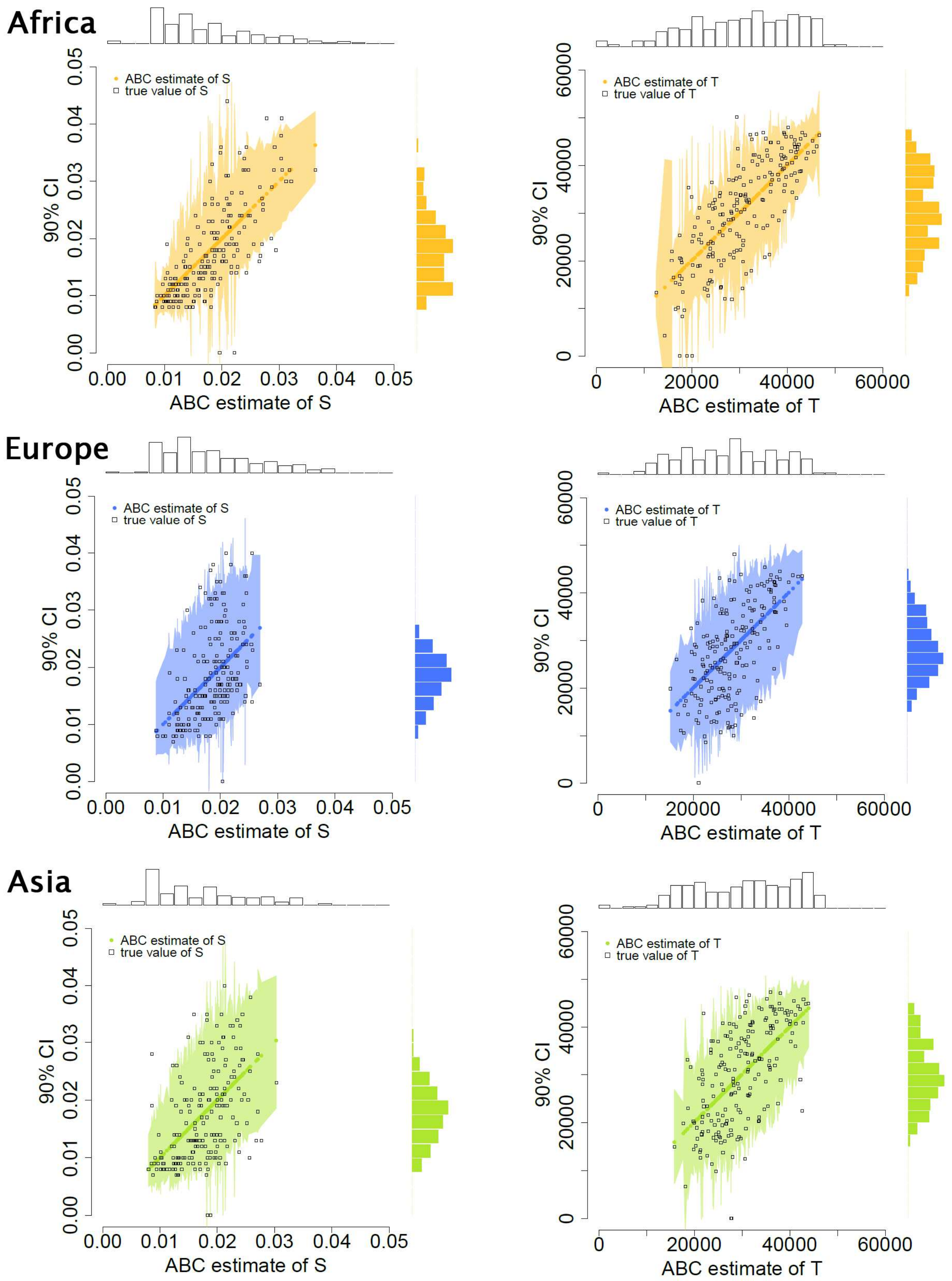


The ABC estimates are those shown in Supplementary Fig. S3. Range of 90% CIs (credible intervals, y-axis) plotted against the ABC estimations (*x*-axis). See also the legend of Supplementary Fig. S6.

**Figure S8. VEP annotations**

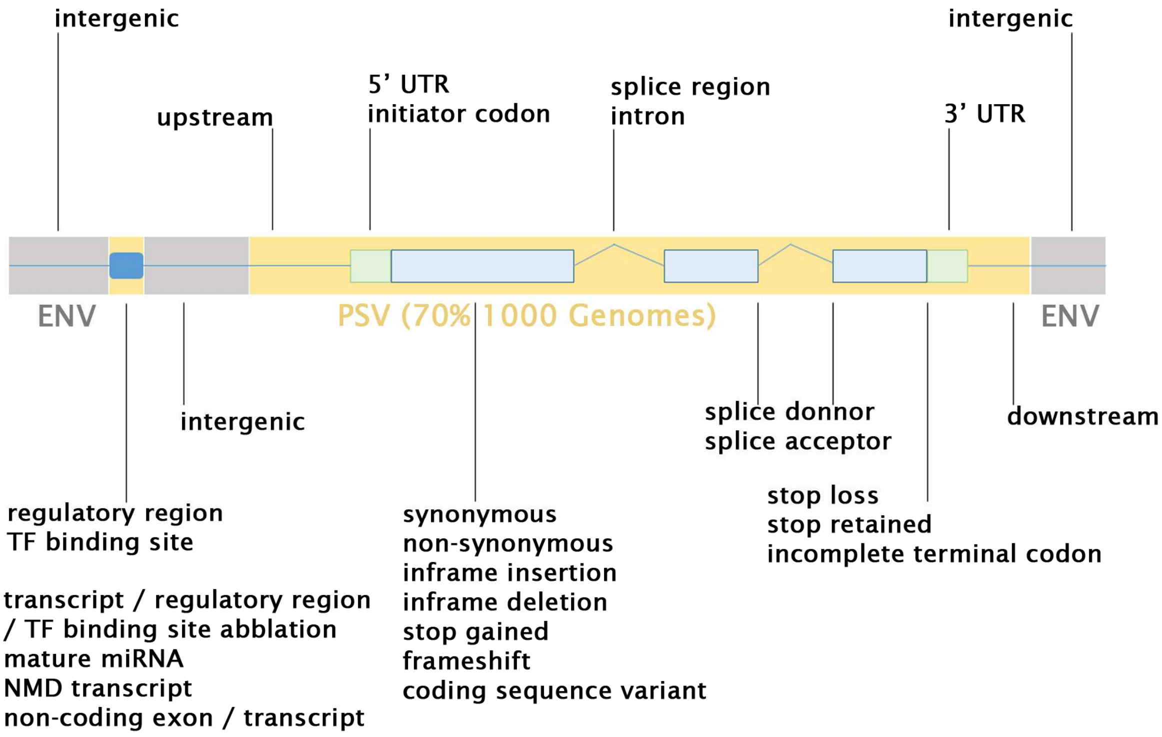


VEP annotation used to define the PSVs and ENVs in the 1000G populations.

**Figure S9. Empirical ORs in each 1000G population**

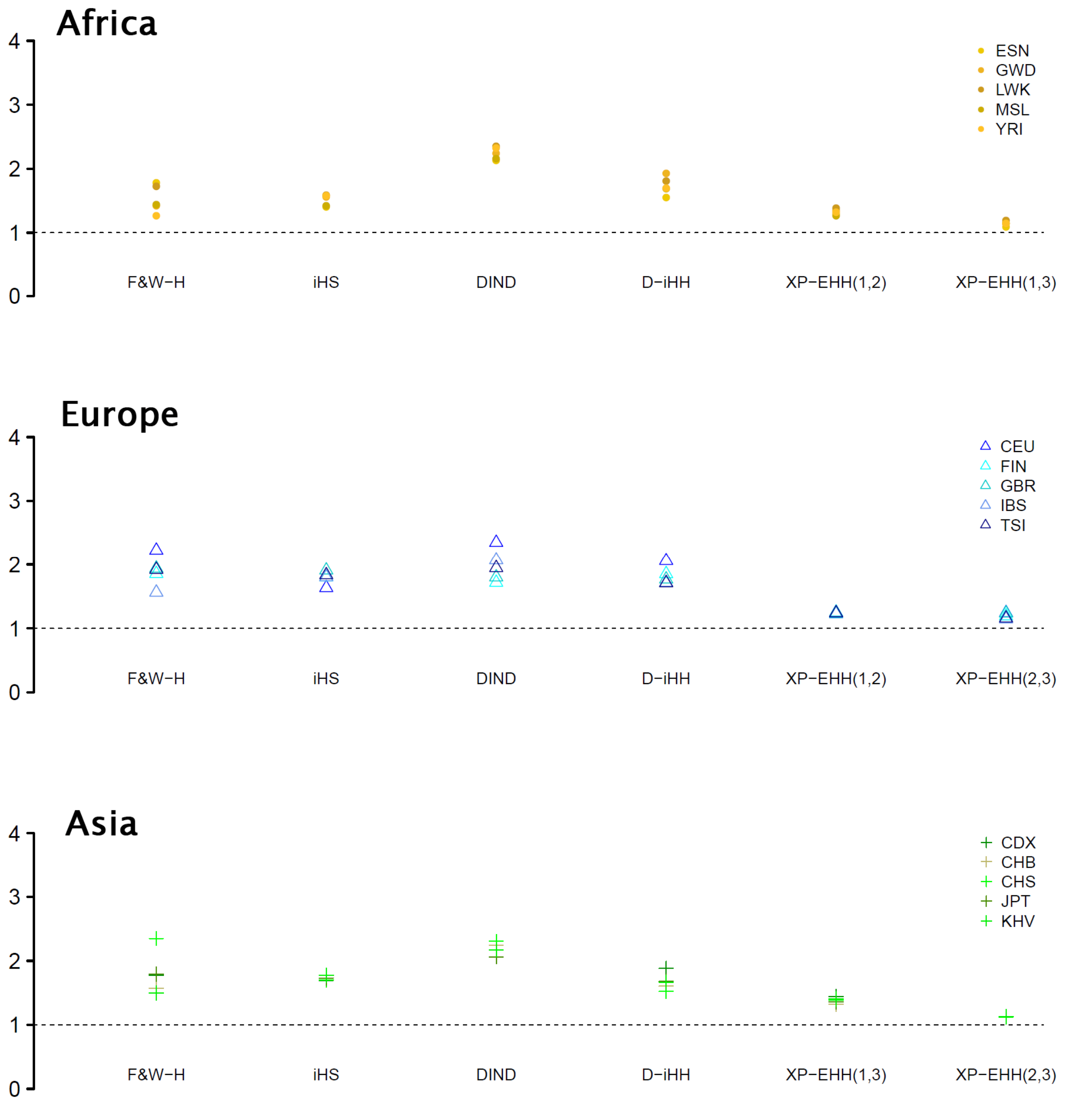


(**A**) ORs computed for each neutrality statistic and corrected for genomic variation in coverage, mutation and recombination rates (Online Methods).

**Figure S10. PCA using simulated and empirical ORs**

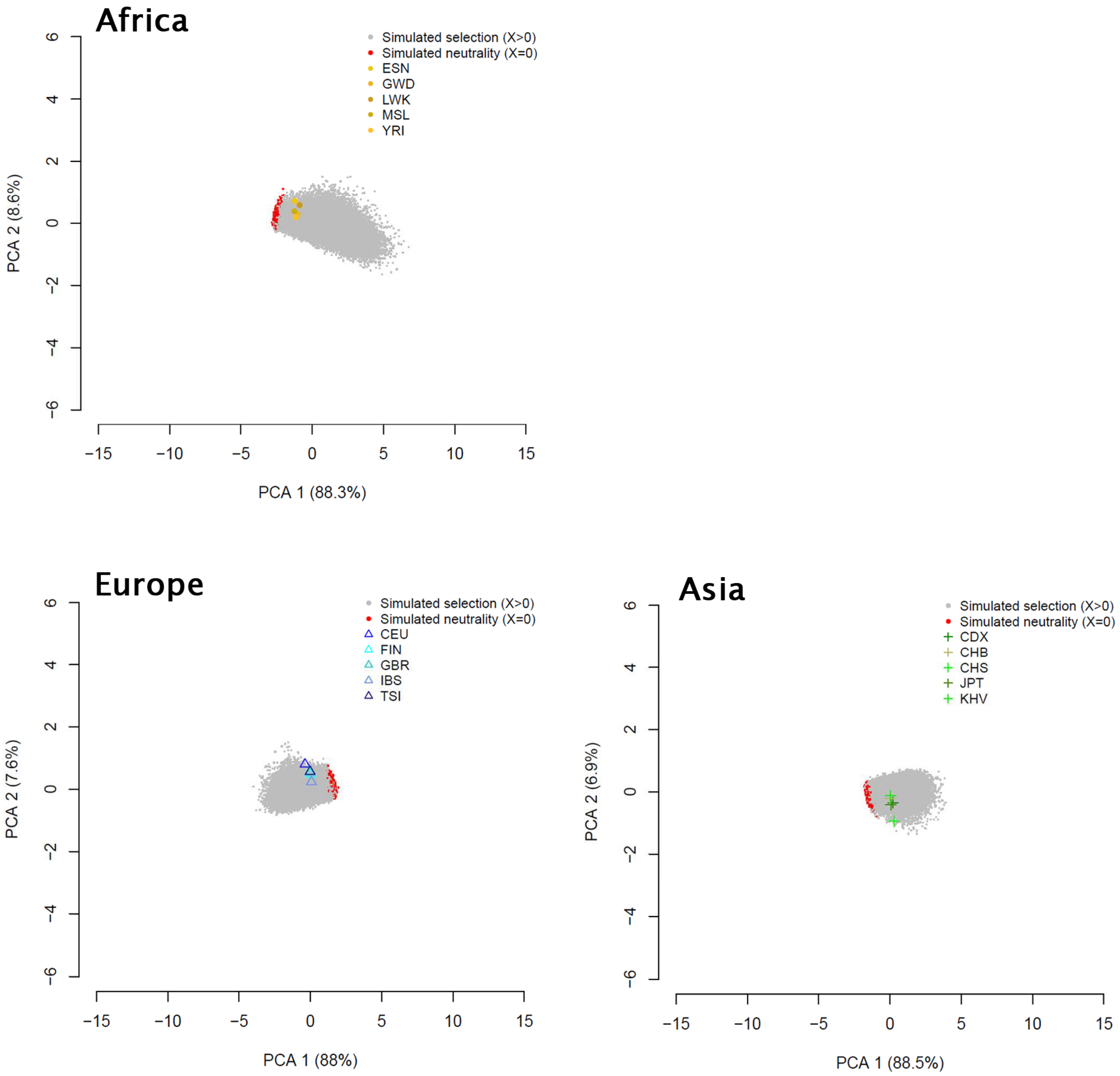


PCA using the simulated ORs used to estimate the selection parameters in each 1000G population (Fig. 3). The empirical ORs observed in African, European and Asian 1000G populations are indicated in corresponding colors. For convenience, we displayed only a small fraction of the $10^5$ simulations performed. These results indicate that the simulated ORs used for the estimations are consistent with the empirical ORs observed in the 1000G populations.

**Figure S11. No correlation between *X* and *S* or *T* estimated in 1000G populations**

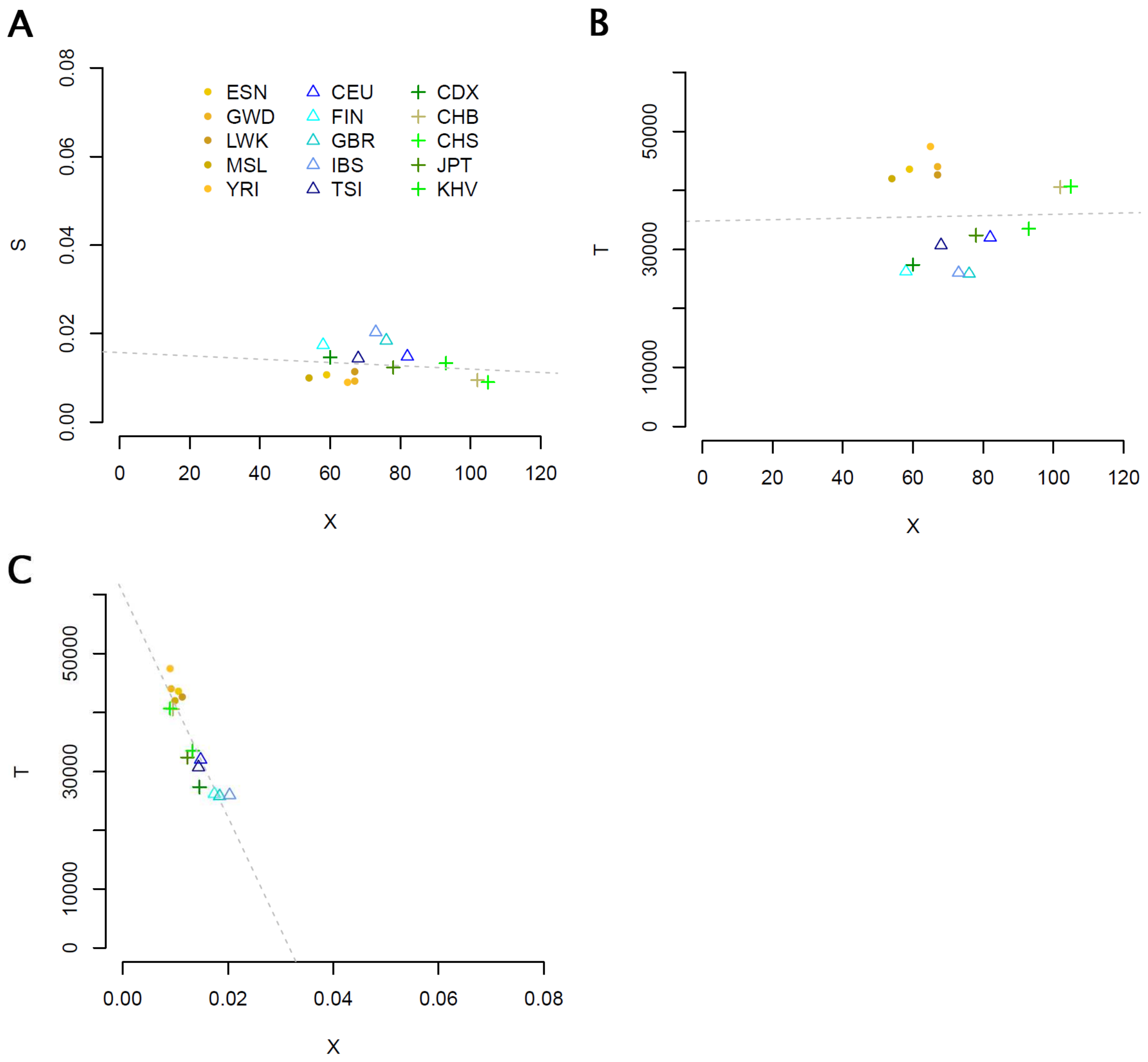


The ABC estimates are those shown in Fig. 3. (**A-B**) No significant correlation between the estimations of *X* and the estimations of *S* or *T*. (**C**) Significant correlation between the estimations of *S* and *T* due to expected correlations in the model itself (see the Supplementary Fig. S5). Regression lines are indicated in grey.

**Figure S12. Posterior predictive checks of the 1000G estimations**

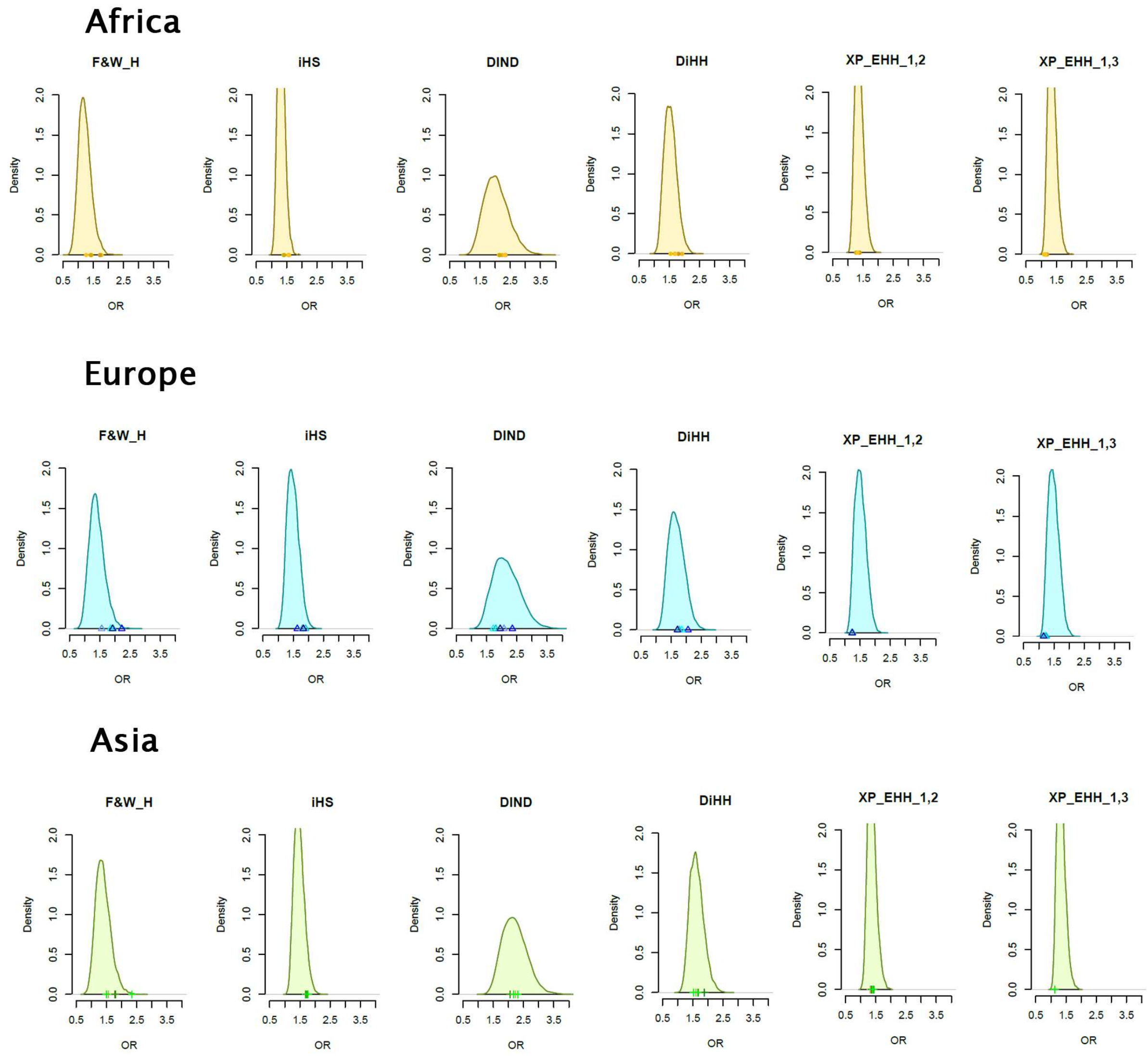


We tested the consistency of the ABC estimations shown in Fig. 3 by performing posterior predictive checks, i.e, simulating summary statistics under the estimated model and comparing them to the empirical summary statistics. We simulated WGS data using parameter values randomly drawn within the CIs of the estimations of *X*, *S* and *T*. Overall, these results show that the re-simulated ORs well match with the empirical 1000G ORs, confirming that our estimations are not biased by empirical data incompatible with the simulated model.

**Figure S13. Estimations using different combinations of neutrality statistics**

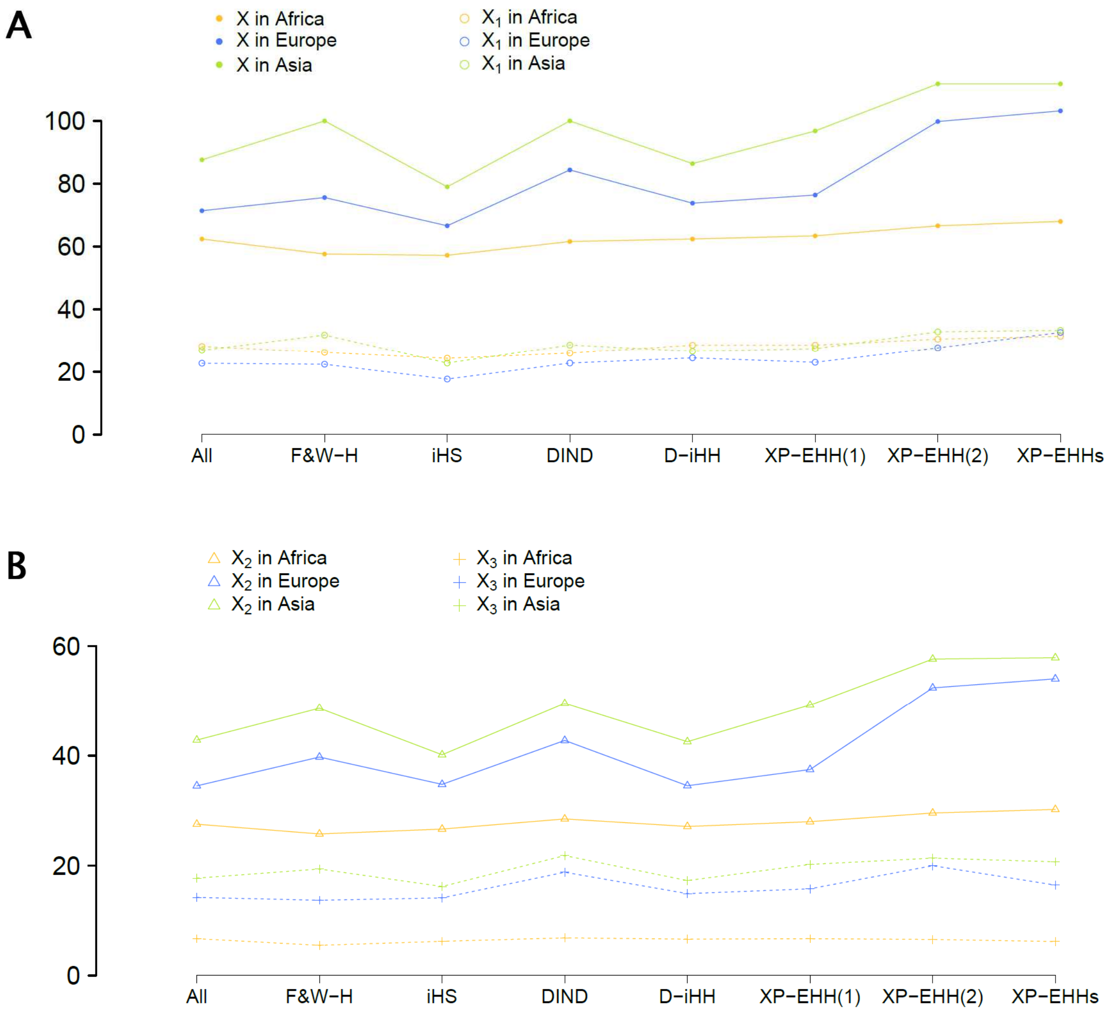


We tested the influence of each summary statistics on the ABC estimations shown in Fig. 3 by performing various sets of new estimations using 5 ORs only, i.e., excluding one summary statistic each time. (**A-B**) Mean numbers of selective sweeps computed across populations of the same continent. 'All' refers to the estimations performed using all summary statistics. Other labels refer to the estimations performed after removing the indicated statistics. XP-EHHs refers to the removal of the two interpopulation statistics (to check potential conflicts between intra and inter population information).

**Figure S14. Influence of the assumptions on the ABC estimations of *X***

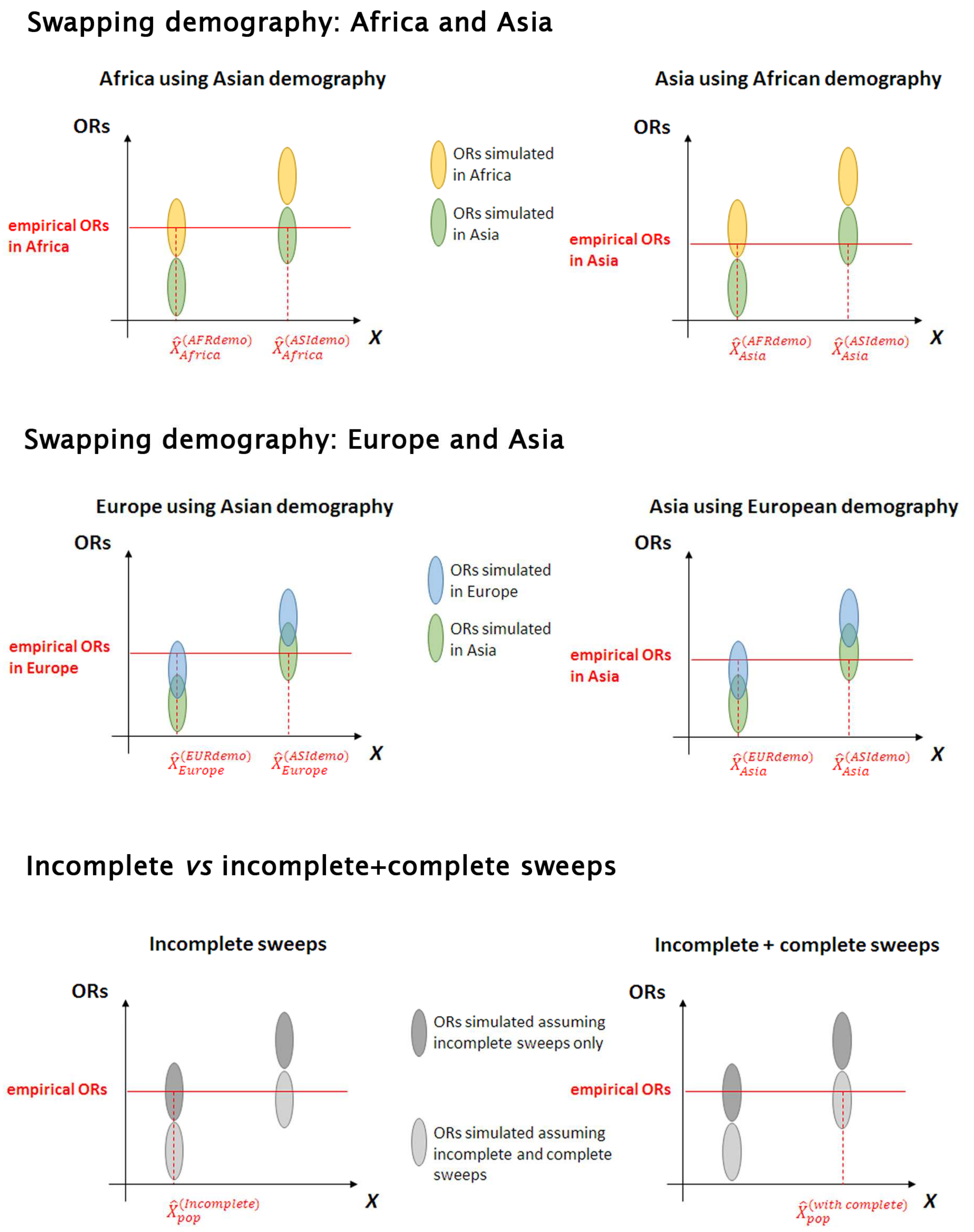


In this explanatory figure, we draw fake distributions of the simulated ORs (y-axis) as represented in Fig. 1 (the differences between ORs are exacerbated for convenience). The $\hat{X}$ (x-axis) represent the estimations of $X$ that can be obtained under various assumptions. The subscripts indicate the population in which the estimations are performed. The superscripts (in brackets) indicate the assumption used, e.g., "AFRdemo" stands for "African demography assumed".

**Figure S15. Estimations using a "wrong" demography: Africa and Asia**

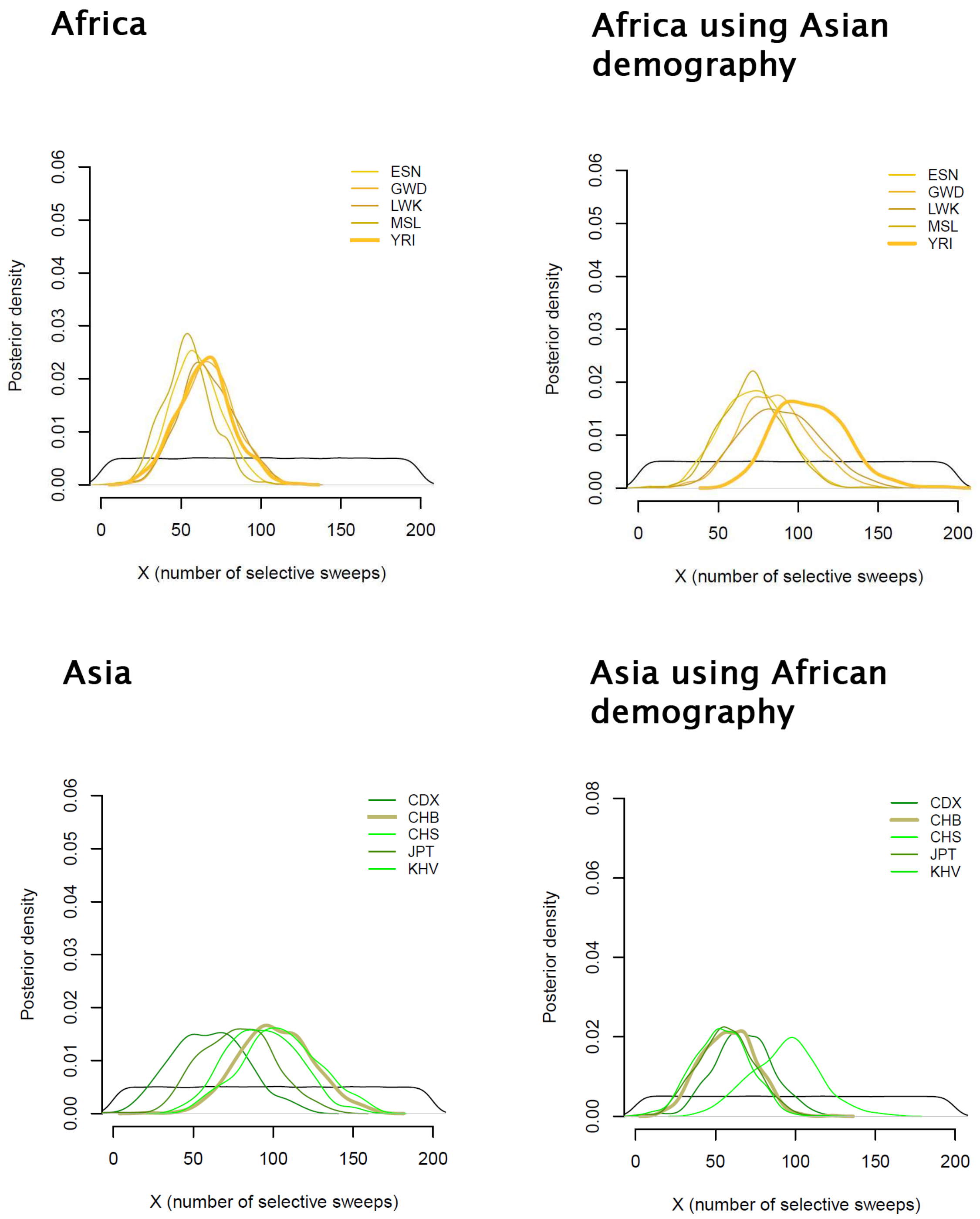


The simulations used to estimate the number of sweeps in African and Asian populations are those used in Fig. 3A-C. The punctual estimates and CIs obtained in each population can be found in Supplementary Table S2. (left panels) Estimations shown in Fig. 3A-C are reported here for comparison. (right panels) Estimations performed by swapping empirical and simulated ORs, i.e., analyzing the African data using the Asian demographic model and inversely.

**Figure S16. Estimations using a “wrong” demography: Europe and Asia**

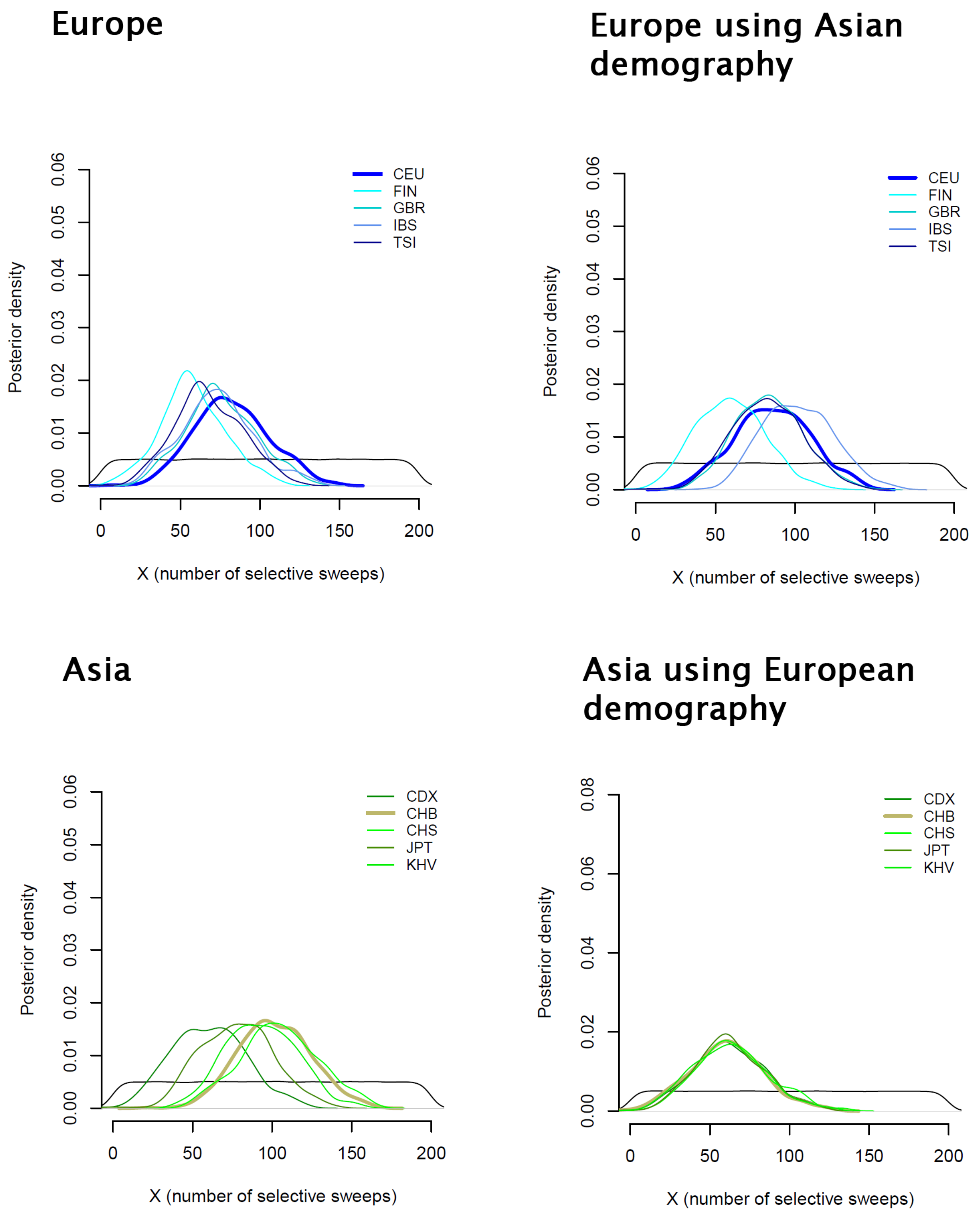


The simulations used to estimate the number of sweeps in European and Asian populations are those used in Fig. 3B-C. The punctual estimates and CIs obtained in each population can be found in Supplementary Table S2. (left panels) Estimations shown in Fig. 3B-C are reported here for comparison. (right panels) Estimations performed by swapping empirical and simulated ORs, i.e., analyzing the European data using the Asian demographic model and inversely.

**Figure S17. Differences in $X$ values due to the bottleneck intensity assumed in the ABC model**

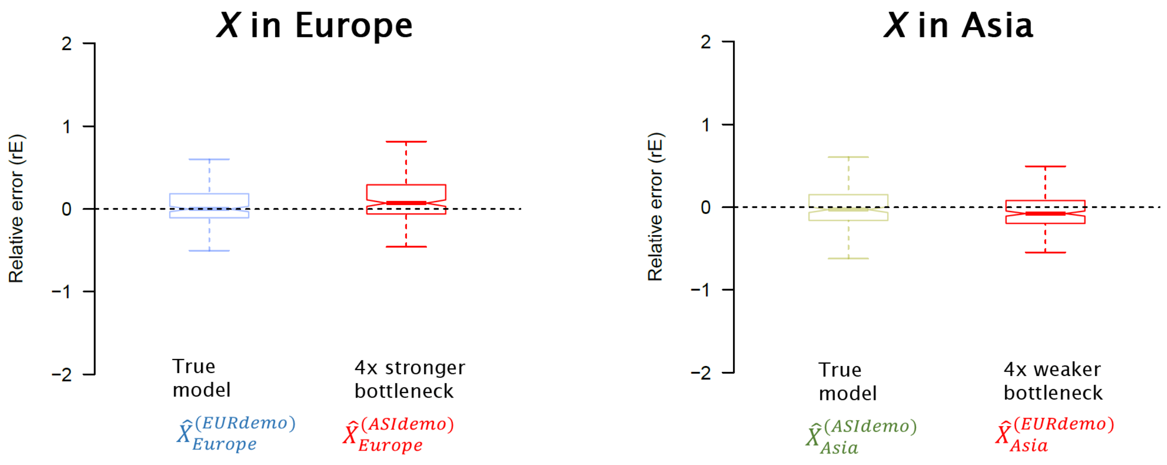


The $\hat{X}$ (x-axis) represent the estimations of $X$ obtained under various assumptions. The subscripts indicate the population in which the estimations are performed. The superscripts (in brackets) indicate the demographic model used, e.g., “EURdemo” stands for “European demography assumed”. (Left hand side) We simulated 200 new WGS European data and analyze them setting (i) a European demography ($\hat{X}^{(EURdemo)}_{Europe}$ labelled “True model”) and (ii) an Asian demography i.e., a four times stronger bottleneck ($\hat{X}^{(ASIdemo)}_{Europe}$ labelled “4x stronger bottleneck”). (Right hand side) We simulated 200 new WGS Asian data and analyze them setting (i) an Asian demography ($\hat{X}^{(ASIdemo)}_{Asia}$ labelled “True model”) and (ii) a European demography i.e., a four times weaker bottleneck ( $\hat{X}^{(EURdemo)}_{Asia}$ labelled “4x weaker bottleneck”).

**Figure S18. Modifying recombination rates, PSVs annotations and OR computations**

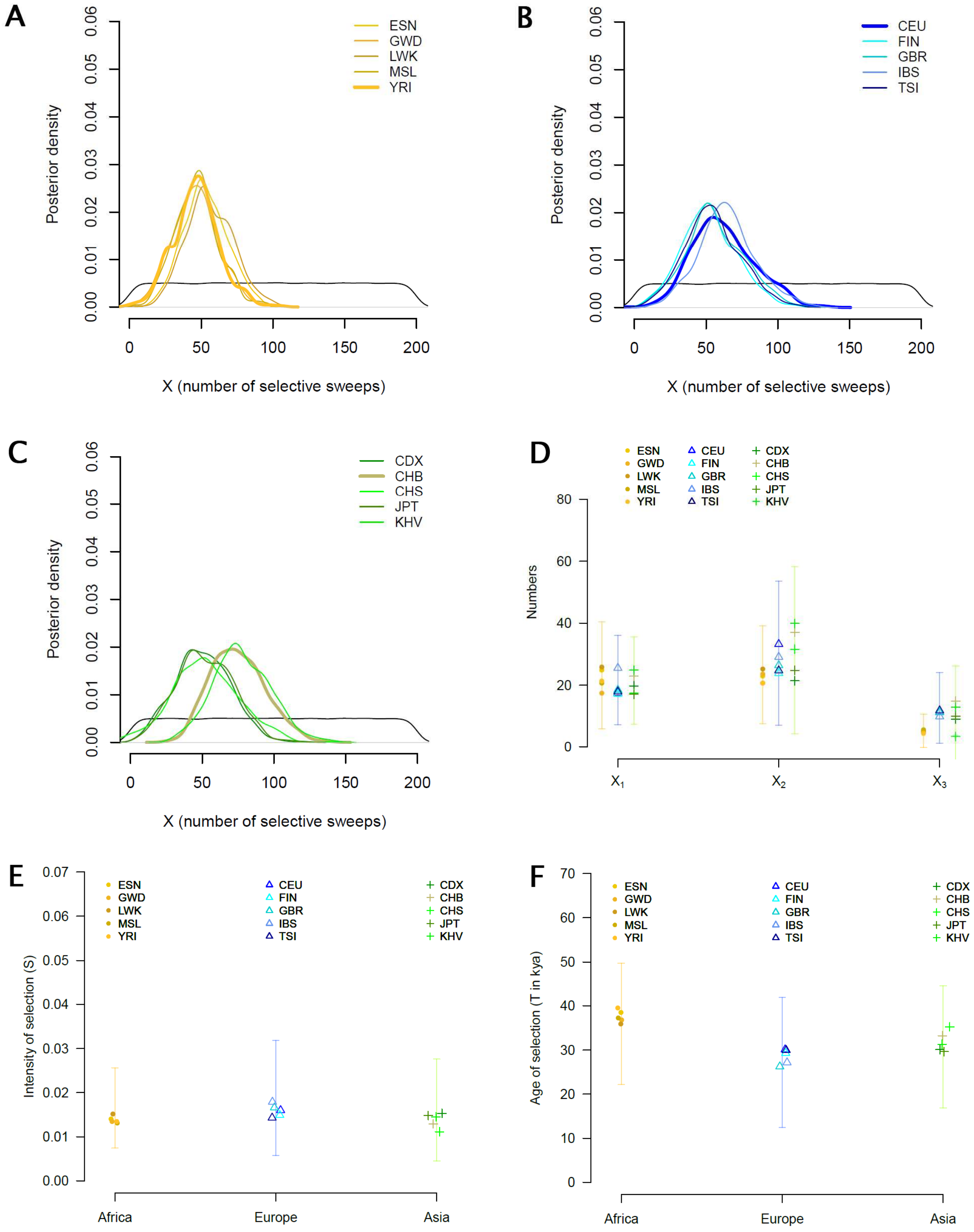


In Fig. 3, we simulated similar recombination rates between ENVs and PSVs (drawings in recombination maps), annotated the selection signals found in ENVs as PSVs and used empirical ORs averaged across chromosomes (Online Methods). In this figure, we set a 10% higher recombination rate in ENVs than in PSVs, excluded selection signal found in ENVs and used empirical ORs computed merging all chromosomes together (empirical ORs are also corrected for genomic variation in coverage, recombination and mutation rates, Online Methods). See the legend of Fig. 3.

**Figure S19. Estimations of *X* including complete sweeps**

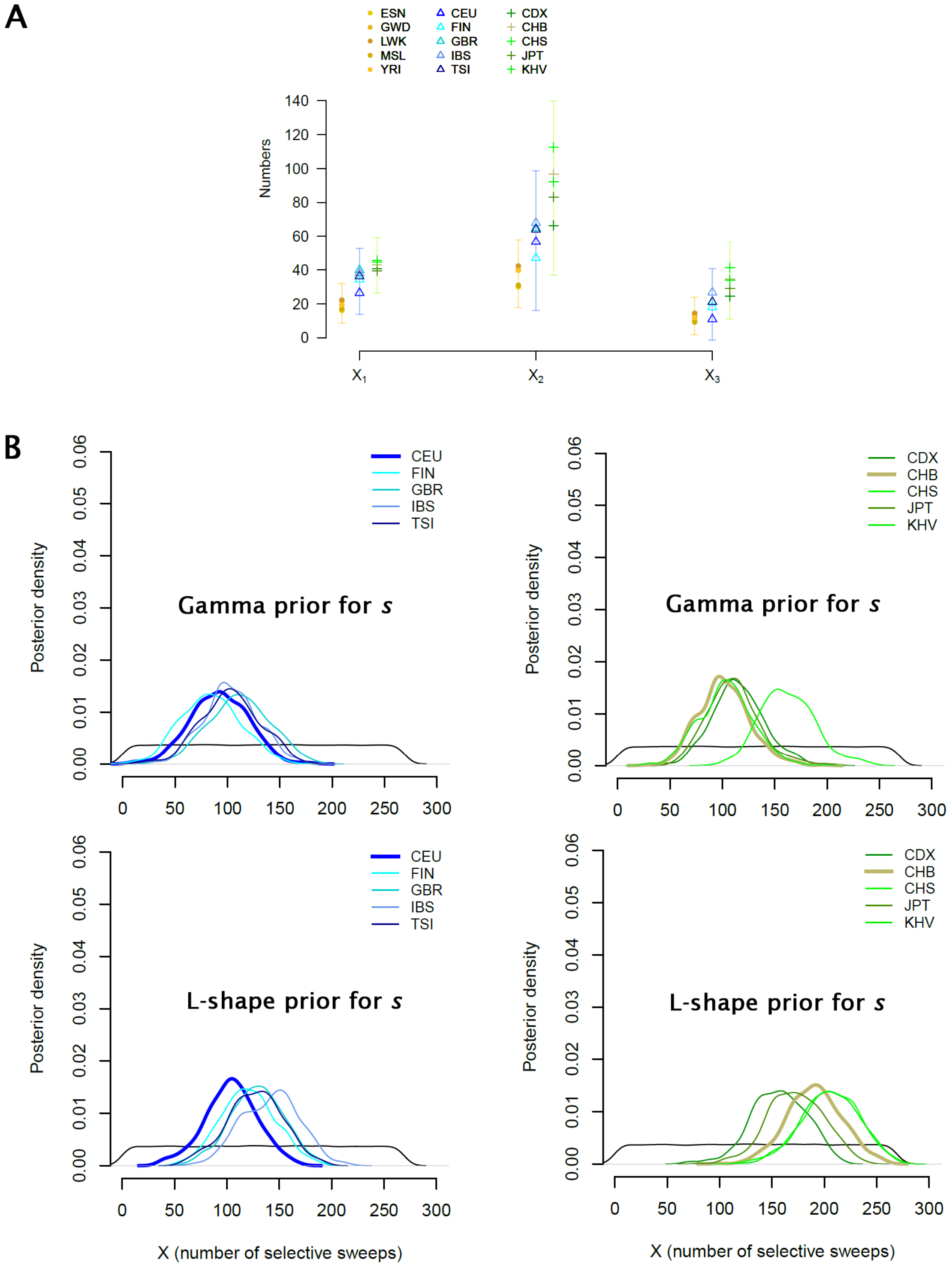


ABC estimations using current selected allele frequencies ranged from 0.2 to 1. (**A**) ABC estimations of $X_1$, $X_2$ and $X_3$, performed using an equal mix of a Gamma and a L-shape prior for *s* (this correspond to the situation shown in Fig. 4). (**B**) ABC estimations of *X* using a Gamma prior (60% of $s \leq 0;01$) and a L-shape prior (90% of $s \leq 0;01$) for *s*. The punctual estimates and CIs can be found in Supplementary Table S2.

**Figure S20. Sharing of selection signals within and between continents**

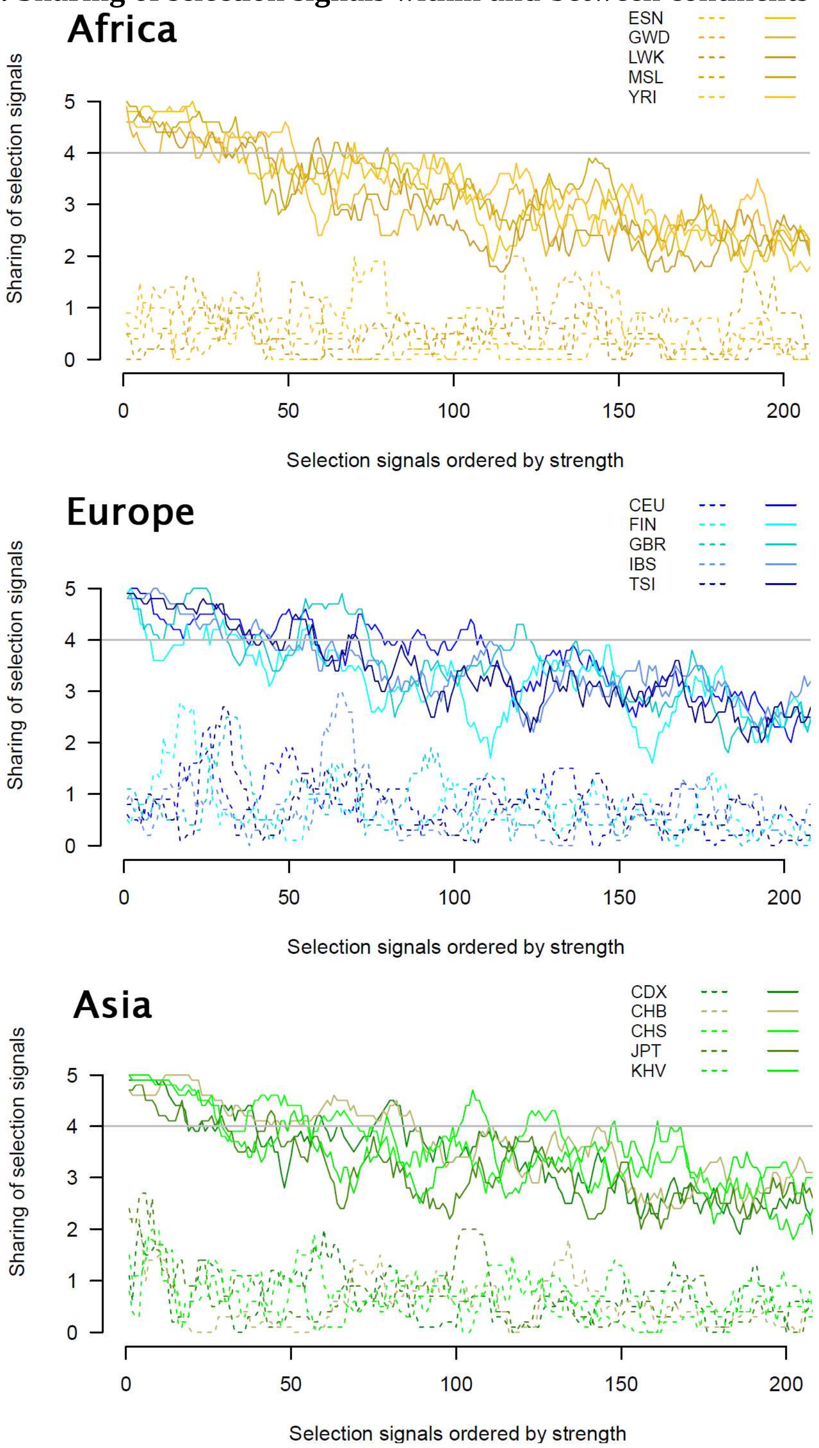


Solid and dashed lines indicate the sharing of selection signals within and between continents. In each population, genomic regions are sorted by signal strength, assessed as the maximal selection score (CSS, Online Methods) found in a region (*x* axis, descendent order). For each population and genomic region enriched in candidate SNPs, we computed an overlap score, i.e., the number of populations for which the same enriched region was identified (detailed results are reported in Supplementary Table S3). With three continents and five populations per continent, the overlap scores computed between continents ranges from 0 (the genomic region was never found enriched in another continent) to 10. The overlap scores computed within each continent ranges from 1 (genomic regions never found to be enriched are not considered) to 5 (the genomic region was found enriched in all populations). We buffered variations by using moving averages of 10 candidate regions.

**Figure S21. Two genomic regions enriched in candidate SNPs**

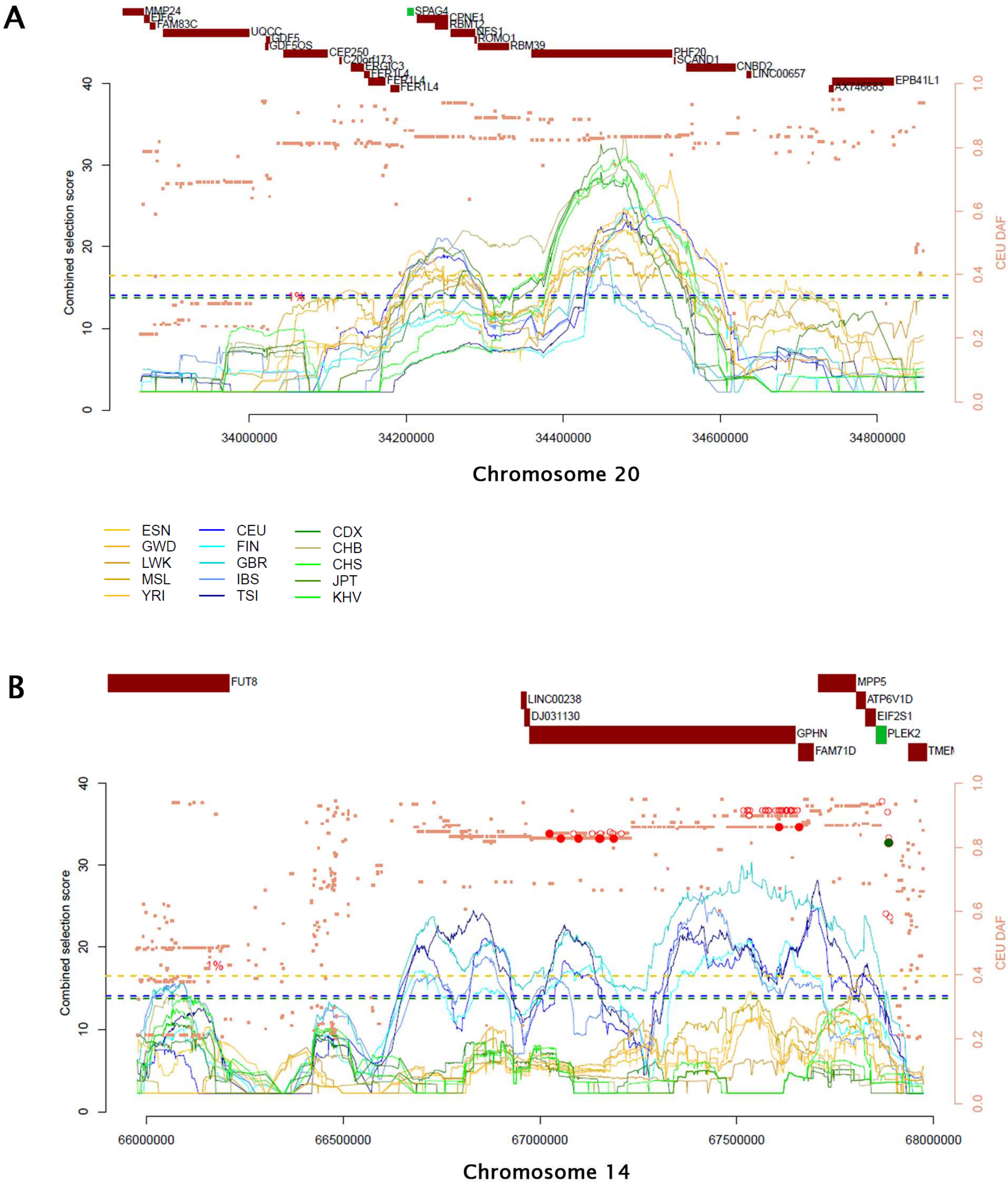


Colored lines are the combined scores of positive selection (CSS, Online Methods), which assess the enrichment of candidate SNPs 100kb around each SNP in each population (dashed lines are the corresponding significance threshold, $P<0.01$). (**A**) A 400kb region significantly enriched in candidate SNPs in all populations (Supplementary Table S3) that encompasses the *SPAG4* gene (in green) associated with impaired sperm motility and infertility in drosophila. (**B**) A ~1 Mb region enriched in candidate SNPs in all European populations (Supplementary Table S3) that encompasses the *PLEK2* gene (in green). Overexpression of *PLEK2* decreases the survival probability in UV-induced melanoma patients (Supplementary Text). Only the allele(s) downregulating *PLEK2* in skin exposed to sun (red dots with the most significant eQTL indicated in green, $P=1.4\times10^{-5}$, GTEx database) harbor signatures indicative of positive selection (closed circles also show genome-wide significant signals of selection revealed in CEU by mean of iHS, DIND and ΔiHH tested separately). Note, the pedigree-based recombination rate (Rutgers maps v3) in PLEK2 is equal to 0.53cM/Mb (Matise, et al. 2007), a value similar to the recombination rate in the LCT region (O'Reilly, et al. 2008).

**Table S1. 1000 Genome phase 3 populations analyzed**

| Names[a] | Populations | Sample sizes |
|---|---|---|
| **African ancestry** | | |
| ESN | Esan | 99 |
| GWD | Gambian | 113 |
| LWK | Luhya | 99 |
| MSL | Mende | 85 |
| YRI | Yoruba | 108 |
| **East Asian ancestry** | | |
| CDX | Dai Chinese | 93 |
| CHB | Han Chinese | 103 |
| CHS | Southern Han Chinese | 105 |
| JPT | Japanese | 104 |
| KHV | Kinh | 99 |
| **European ancestry** | | |
| CEU | CEPH (Utah residents) | 99 |
| GBR | British | 91 |
| FIN | Finnish | 99 |
| IBS | Spanish | 107 |
| TSI | Tuscan in Italy | 107 |
| Total | | 1511 |

[a]ESN (Esan in Nigeria), GWD (Gambian in Western Division, Mandinka), LWK (Luhya in Webuye, Kenya), MSL (Mende in Sierra Leone) and YRI (Yoruba in Ibadan, Nigeria), CEU (Utah residents), GBR (British in England and Scotland), FIN (Finnish in Finland), IBS (Iberian Populations in Spain) and TSI (Tuscan in Italy), CDX (Chinese Dai in Xishuangbanna, China), CHB (Han Chinese in Beijing, China), CHS (Southern Han Chinese, China), JPT (Japanese in Tokyo, Japan) and KHV (Kinh in Ho Chi Minh City, Vietnam).

**Table S2**. **Parameters estimated in each 1000G population (additional Excel file)**

**Table S3**. **Genomic regions significantly enriched in candidate SNPs of selection (additional Excel file)**

### Supplementary text

#### Genomic regions with the highest enrichment in candidate SNPs of selection

The genomic regions with significant enrichment in candidate SNPs (Supplementary Table S3) confirms various adaptations to recent selective pressures including pathogens, nutrition, and climate. We retrieved the well-documented examples of positive selection associated with lactase persistence in adulthood (*LCT*) in northern Europe and the genomic region of *EDAR* previously found to be under selection in Asia. The *TLR5* region also detected under positive selection in the same 1000G populations (Grossman, et al. 2013) has been found to be among the 20 most enriched regions in all African populations except one. We retrieved other examples of well-described candidate regions for selection (Vitti, et al. 2013; Jeong and Di Rienzo 2014; Fan, et al. 2016), e.g., *APOL1* (resistance to human trypanosomes causing "African sleeping sickness"), *SLC45A2* (lighter pigmentation in Europe), *HERC2* (rs12913832 associated light skin pigmentation in Europeans (Key, et al. 2016)), *SPAG4* corresponding to the strongest iHS signal previously found in Europeans (Voight, et al. 2006), the *TLR1-6-10* cluster in Europe (Quach, et al. 2016), *BBX* (Grossman, et al. 2013), *HERC1* (Grossman, et al. 2013) and the *ADH* cluster (alcohol dehydrogenase) in Asians (Barreiro, et al. 2008) (Supplementary Table S3). However, because we did not consider neutrality statistics potentially inflated by background selection, such as the fixation index $F_{ST}$, some genomic regions previously detected on the basis of larges differences in allele frequencies between populations only are not replicated in this study, such as the *DARC* region (reduced susceptibility to malaria in Africa) (Hamblin and Di Rienzo 2000).

#### Evidences supporting positive selection in the PLEK2 region

One ~1Mb region significantly enriched in candidate SNPs was detected in European populations only (Supplementary Table S3). This region encompasses functional variants controlling the expression of the *PLEK2* gene in skin cells exposed to the sun, i.e., eQTLs in the GTEx database (Ardlie, et al. 2015) (no eQTL found in skin not exposed to sun (Ardlie, et al. 2015)). *PLEK2* overexpression causes large lamellipodia (Hu, et al. 1999), is characteristic of disseminated tumor cells (Naume, et al. 2007), and has been detected in analyses of the blood of ~80% of melanoma patients. *PLEK2* may serve as a biomarker for the disease (Luo, et al. 2011). In addition, the survival probability was found diminished in patients with the highest *PLEK2* overexpression (https://www.proteinatlas.org/ENSG00000100558-PLEK2/pathology/tissue/melanoma). At the protein level, expression has been found higher in melanoma cells than in normal tissue (Uhlen, et al. 2015).

In this study, we found that alleles downregulating *PLEK2* harbor signatures indicative of positive selection. Interestingly, these variants which affect sensitivity to UV-induced melanoma appear to have been under selection in European populations, while this selection signal was not found in Africa and Asia (Supplementary Table S3). Prevalence of UV-induced melanoma is more than 20 times more common in European Americans than in African Americans (the lifetime risk of getting melanoma is 2.5% and 0.1% in European and African Americans respectively, the American Cancer Society). Finally, in current Europeans, the level of *PLEK2* expression in skin and the risk of developing UV-induced melanoma both increase with age (Glass, et al. 2013) (average of age of patients is ~63 years old in US but melanoma is not uncommon among those younger than 30, the American Cancer Society). These observations are consistent with the role of natural selection in the maintenance of low *PLEK2* expression in early stages of life (during the reproductive period). In main text, we speculated that alleles downregulating *PLEK2* could have been favored by selection during the late Pleistocene climatic warming (~20 to ~10kya). This speculative model needs to be further tested by formally estimating the age of selection in the region (selection signal of ~1-1.2Mb suggests a recent onset of selection).
